# SNARE Complex Alters the Interactions of the Ca^2+^ sensor Synaptotagmin 1 with Lipid Bilayers

**DOI:** 10.1101/2020.06.19.161752

**Authors:** Maria Bykhovskaia

## Abstract

Release of neuronal transmitters from nerve terminals is triggered by the molecular Ca^2+^ sensor Synaptotagmin 1 (Syt1). Syt1 is a transmembrane protein attached to the synaptic vesicle (SV), and its cytosolic region comprises two domains, C2A and C2B, which are thought to penetrate into lipid bilayers upon Ca^2+^ binding. Prior to fusion, SVs become attached to the presynaptic membrane (PM) by the four-helical SNARE complex, which binds the C2B domain of Syt1. To understand how the interactions of Syt1 with lipid bilayers and the SNARE complex trigger fusion, we performed molecular dynamics (MD) simulations at a microsecond scale. The MD simulations showed that the C2AB tandem of Syt1 can either bridge SV and PM or immerse into PM, and that the latter configuration is more favorable energetically. Surprisingly, C2 domains did not cooperate in penetrating into PM, but instead mutually hindered the lipid penetration. To test whether the interaction of Syt1 with lipid bilayers could be affected by the C2B-SNARE attachment, we performed systematic conformational analysis of the Syt1-SNARE complex. Notably, we found that the C2B-SNARE interface precludes the coupling of C2 domains of Syt1 and promotes the immersion of both domains into the PM bilayer. Subsequently, we simulated this pre-fusion protein complex between lipid bilayers imitating PM and SV and found that the immersion of Syt1 into the PM bilayer within this complex promotes PM curvature and leads to lipid merging. Altogether, our MD simulations elucidated the role of the Syt1-SNARE interactions in the fusion process and produced the dynamic all-atom model of the prefusion protein-lipid complex.

**Statement of Significance:** Neuronal transmitters are packed in synaptic vesicles (SVs) and released by fusion of SVs with the presynaptic membrane (PM). SVs are attached to PM by the SNARE protein complex, and fusion is triggered by the Ca^2+^ sensor Synaptotagmin 1 (Syt1). Although Syt1 and SNARE proteins have been extensively studied, it is not yet fully understood how the interactions of Syt1 with lipids and the SNARE complex induce fusion. To address this fundamental problem, we took advantage of Anton2 supercomputer, a unique computational environment, which enables simulating the dynamics of molecular systems at a scale of microseconds. Our simulations produced a dynamic all-atom model of the prefusion protein-lipid complex and demonstrated *in silico* how the Syt1-SNARE complex triggers fusion.

## Introduction

Neuronal transmitters are released from nerve terminals in response to Ca^2+^ influx. Calcium binds Synaptotagmin 1 (Syt1), a synaptic vesicle (SV) protein which drives the fusion of SVs with the plasma membrane (PM) and triggers rapid transmitter release [1].

Syt1 comprises two Ca^2+^ binding domains, C2A and C2B, connected by a flexible linker and attached to SV by a transmembrane helix [2]. Each domain has two loops forming a Ca^2+^ binding pocket, and in each pocket Ca^2+^ ions are chelated by five aspartic acids [3-5]. Both domains, C2A [3, 6] and C2B [4] can bind phospholipids, and the binding properties of C2 domains depend on the lipid composition. In particular, the association of the C2B domain with the lipid membrane depends on the presence of phosphatidylinositol 4,5-bisphosphate (PIP_2_) in the bilayer [7].

It is generally agreed that SV-PM fusion depends on the interaction of Syt1 into phospholipids [8], however it is still debated how the protein-lipid occurs at the atomistic level. One possible scenario is that Syt1 bridges PM and SV by inserting the Ca^2+^ bound tips of its C2 domains into the opposing phospholipid bilayers [9, 10]. Alternatively, Syt1 could trigger the fusion by penetrating into PM with the tips of its both domains, promoting PM curvature, and generating the lipid stalk followed by pore opening [11, 12]. It was also proposed that these two mechanisms could be combined and that Syt1 could bridge the membranes via a heteromerization [13, 14]. Finally, different configurations of Syt1 domains could control different modes of transmitter release, such as synchronous, asynchronous, and spontaneous fusion [15].

The attachment between SV and PM is maintained by the protein complex termed SNARE [16, 17]. The SNARE proteins form a coil-coiled four-helical bundle, which consists of the SV-associated protein synaptobrevin (Syb) and the PM-associated proteins syntaxin (Syx) and SNAP25 also known as t-SNARE. Syt1 can interact with t-SNARE proteins, and multiple studies suggest an important role for Syt1-SNARE interactions during fusion [18-23]. However, other studies have argued against this possibility [24, 25], and it is still debated how the Syt1-SNARE complex is formed *in vivo* and what is the role of the Syt1-SNARE interactions in the fusion process [26-32].

Fusion is regulated by the cytosolic protein Complexin (Cpx), which attaches to the SNARE complex [33], forming a five helical SNARE-Cpx bundle. Cpx promotes Ca^2+^-dependent fusion [34-36], and several structural [37, 38], genetic [39] and biochemical [40, 41] studies suggest that Cpx may directly interact with Syt1 on the SNARE bundle (but see also [42]).

To elucidate the atomistic detail of Syt1 interations with the SNARE-Cpx complex and lipid bilayers, we performed all-atom molecular dynamics (MD) simulations at a microsecond scale. We took advantage of unique capabilities of the specialized Anton supercomputer designed for MD simulations [43, 44], which enabled a breakthrough in simulating the protein dynamics at a time scale of microseconds [45, 46]. We have recently employed these computational tools to investigate the conformational dynamics of Syt1 in the solution [47, 48]. Here, we employed this approach to elucidate how Syt1 interactions with PM, SV, and the SNARE-Cpx complex could drive synaptic fusion.

## Methods

### System setup

All the molecular systems were constructed using Visual Molecular Dynamics Software (VMD, Theoretical and Computational Biophysics Group, NIH Center for Macromolecular Modeling and Bioinformatics, at the Beckman Institute, University of Illinois at Urbana-Champaign). All the simulations were performed in water/ion environment with explicit waters. Potassium and chloride ions were added to neutralize the systems and to yield 150 mM concentration of KCl. Water boxes with added ions were constructed using VMD.

The palmitoyl oleyl phosphatidylcholine (POPC) lipid bilayers were generated using VMD. The initial structure of anionic lipid bilayer containing phosphatidylserine (POPS) and PIP_2_, POPC:POPS:PIP_2_ (75:20:5) [74] was kindly provided by Dr. J. Wereszczynski (Illinois Institute of Technology). In all the systems, the lipid bilayers were positioned in the XY plain.

The initial structures for the isolated C2A [75]) and C2B [76] domains in their Ca^2+^ - bound forms were obtained from crystallography studies (1BYN and 1TJX, respectively in the Protein Data Base). The initial structures of the isolated domains in their Ca^2+^ free forms were obtained by removing Ca^2+^ ions from the Ca^2+^-bound structures. For the initial structure of the Syt1 C2AB tandem, we took a transient state with uncoupled domains from the MD trajectory obtained in our earlier study [48].

To build the initial model of the C2AB-SNARE-Cpx complex (I1), we combined two structures: 5CCG (C2AB-SNARE complex) obtained by crystallography [27] and the SNARE-Cpx complex extracted from our earlier MD simulations [54]. We have superimposed the SNARE bundles in the two models by minimizing their RMSD distances. Subsequently, one of the SNARE bundles (from 5CCG) was removed, and the structure of the resulting Syt1-SNARE-Cpx complex was optimized employing energy minimization. Chelated Ca^2+^ ions were kept within Ca^2+^ binding pockets, as in the original 5CCG structure. To build the structure of the C2B-SNARE-Cpx complex, we have removed the C2A domain (residues 140-270) from the Syt1-SNARE-Cpx complex.

The additional starting models of the Syt1-SNARE-Cpx complex (I3 and I4) were built as follows. The starting configuration for the I3 complex was derived from the 3N1T C2B-SNARE complex [28] combined with the C2AB tandem [33, 48] and the SNARE-Cpx complex [54]. The structures were combined subsequently. First, the C2B domain from C2B-SNARE (3N1T) complex was superimposed with the C2B domain from the C2AB tandem employing the RMSD minimization, and then the C2B module from 3N1T was removed. Subsequently, the SNARE bundle from the resulting C2AB-SNARE complex was superimposed with the SNARE-Cpx complex employing the same method, and the first SNARE bundle was removed. This procedure was followed by the energy minimization of the resulting C2AB-SNARE-Cpx complex.

The starting configuration for the I4 complex was obtained by docking of the C2AB tandem to the SNARE-Cpx complex. We have positioned the SNARE-Cpx complex and the C2AB tandem in such a way that the Syt1 - interacting stretch of SNAP25 (32-40, [58]) was facing the SNARE-interacting surface of the C2B domain [47]. Out of the two possible orientations of the C2B domain, we chose the one with negatively charged residues of Syt1 facing the positively charged residues of SNAP25 and *vice versa*. Subsequently, we performed the Monte-Carlo/Minimization docking procedure [77] for this molecular system employing ZMM software (www.zmmsoft.com [78]).

All the molecular systems, including their sizes and the lengths of respective MD trajectories, are summarized in the Tables S1.

### Molecular dynamics

The MD simulations were performed employing CHARMM36 force field [79] modified to include the parameters for PIP_2_ as described in [74]. The simulations were performed with periodic boundary conditions and Ewald electrostatics in NPT ensemble at 310K.

The heating (20 ps) and equilibration (100 ns) phases were performed employing NAMD [80] Scalable Molecular Dynamics (Theoretical and Computational Biophysics Group, NIH Center for Macromolecular Modeling and Bioinformatics, at the Beckman Institute, University of Illinois at Urbana-Champaign) at XSEDE (Extreme Science and Engineering Discovery Environment) Stampede cluster (TACC). The NAMD simulations were performed with a flexible cell and with a time-step of 1.5 fs, employing Langevin thermostat and Berendsen barostate.

Production runs were performed at Anton2 supercomputer [43, 44] with Desmond software through the MMBioS (National Center for Multiscale Modeling of Biological Systems, Pittsburg Supercomputing Center and D.E. Shaw Research Institute). All the Anton2 simulations were performed in a semi-isotropic regime, with a time-step of 2.5 fs, and employing the multigrator [81] to maintain constant temperature and pressure.

### Analysis and Visualization

The trajectory analysis was performed employing VMD and Vega ZZ (Drug Design Laboratory) software. All the parameters along all the trajectories were computed with a time step of 2.4 ns. The number of VdW contacts between two molecules was computed as the number of the pairs of atoms separated by the distance of less than 3 Å. The distributions of the parameters along MD trajectories were compared using the Kolomogorov-Smirnov (K.-S.) test.

## Results

### The interactions of the C2B domain of Syt1 with phospholipid bilayers and the SNARE complex

We started from simulating the interactions of the C2B domain with lipid bilayers, since these interactions were shown to be critical for the fusion process [8, 49, 50]. First, we simulated the Ca^2+^ -free and Ca^2+^ - bound (Ca^2+^C2B) forms of the C2B domain at homogeneous neutral lipids (POPC). In the initial configuration (Fig. 1S A, Supporting Material), the protein was positioned in such a way that it did not form any Van der Waals (VdW) contacts with the bilayer. This initial configuration was also used in all the subsequent simulations of the isolated C2B domain.

**Figure 1.**
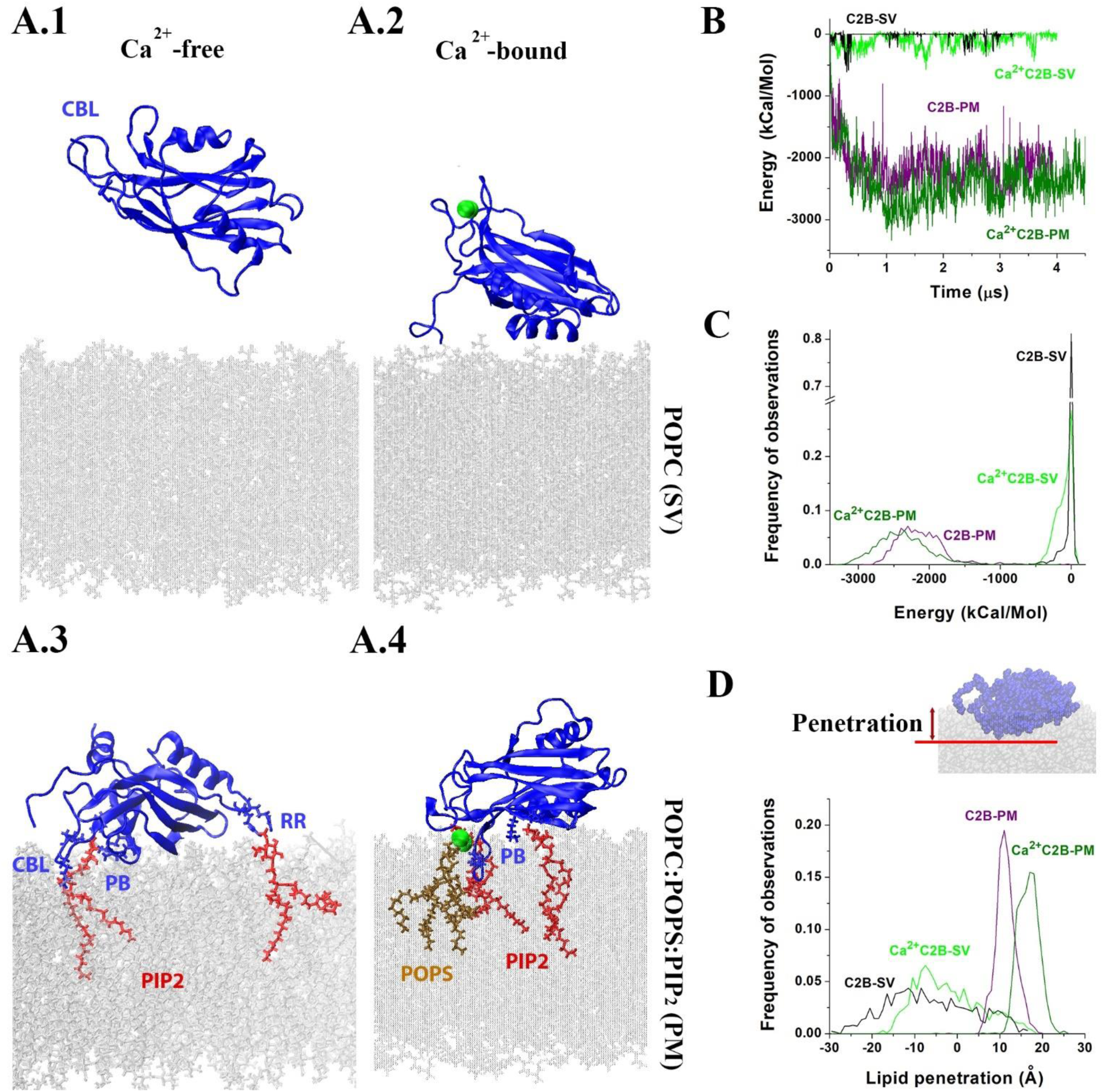
Interactions of the C2B domain with lipid bilayers. **A**. Representative conformational states at the end of respective trajectories showing the attachment of the Ca^2+^ free or Ca^2+^ bound C2B module to the POPC or POPC:POPS:PIP_2_ bilayer. Green spheres depict Ca^2+^ ions. **A**.**1**. The Ca^2+^ free C2B module is detached from the SV bilayer. **A**.**2**. The Ca^2+^C2B module interacts with the SV bilayer via its α-helix at the opposite tip from CBL. **A**.**3**. The C2B module is anchored to PIP_2_ molecule via its PB and CBL motifs, while the RR motif anchors to another PIP_2_ molecule. **A**.**4**. CBL of the Ca^2+^C2B module is immersed into the bilayer, being anchored to a PIP_2_ molecule, while Ca^2+^ ions form coordination bonds with two POPS molecules. In addition, PB is anchored to another PIP_2_ molecule. **B**. The energies of the interactions of C2B and Ca^2+^C2B with POPC and POPC:POPS:PIP_2_ lipid bilayers imitating the SV and PM, respectively. The energies are plotted along the respective MD trajectories. **C**. Energy distributions of the interactions of C2B and Ca^2+^C2B with lipid bilayers imitating SV and PM. The stronger interactions correspond to more negative values. **D**. Lipid penetration of C2B and Ca^2+^C2B. More positive values correspond to deeper immersion of the protein into lipids. Ca^2+^ binding significantly promotes the penetration (p<0.01 per K-S test).

The 3.2 μs production run of the Ca^2+^-free C2B module showed that the protein was largely detached from the bilayer over the entire length of the MD trajectory, making infrequent contacts with lipids (Fig. S1 B.1, B.2; Fig. 1 A.1). The analysis of energy components demonstrated that electrostatic energy prevails in the C2B-lipid interactions, and also that the energy of C2B-ion interactions drastically exceeds the energy of C2B-lipid interactions (Fig. S1, B.3). This result suggests that the Ca^2+^-free C2B module has a preference for water/ion environment versus lipids, and it explains the limited number of protein-lipid contacts observed over the course of the MD trajectory (Fig. S1 B.1).

In contrast, the Ca^2+^C2B domain largely interacted with the bilayer over the course of the 4.0 μs trajectory, and these interactions were largely maintained by its alpha helix and the adjacent beta sheets (residues 380-410), as well as by its C2AB linker (Fig. S1 C.1, C.2; Fig. 1 A.2). The analysis of energy components showed that the energies of Ca^2+^C2B-ion interactions were shifted towards more positive values when compared to the Ca^2+^-free module (Fig. S1 D, top), possibly due to the reduced electrostatic attraction to K^+^ ions. Respectively, the energies of Ca^2+^C2B-lipid interactions were shifted towards more negative values (Fig.1S D, bottom), in agreement with consistent protein-lipid contacts observed for the Ca^2+^C2B module (Fig. 1S, C) but not for its Ca^2+^-free form.

The interactions of the C2B module with neutral phospholipids described by these simulations would likely define the dynamics of the attachment of the C2B domain to SV, since SVs are predominantly composed out of neutral lipids [51, 52]. Our results suggest that the Ca^2+^-free C2B domain would likely float in a cytosol being detached from SV, while Ca^2+^ binding would promote a week C2B-SV association. We next asked how would the C2B module interact with PM, which has a high content of anionic lipids and also incorporates PIP_2_ [8, 53], which is critical for the attachment of Syt1 [7, 10, 25]. To imitate the PM composition, we used the atomic model of the membrane patch containing anionic lipids (POPS, 20%) and PIP_2_ (5%).

The simulations demonstrated that both Ca^2+^ -free and Ca^2+^-bound forms of the C2B domain rapidly attached to the POPC:POPS:PIP_2_ (75:20:5) bilayer and stayed attached over the entire course of respective trajectories (Fig. S2 A, blue; Fig. 1 A.3). The analysis of energy components demonstrated the C2B-lipid attraction was largely defined by the interactions of the C2B module with PIP_2_ (Fig. S2 B). The interactions with PIP_2_ were maintained by the three sites of the C2B domain (Fig. S2 C): 1) the polybasic stretch (PB, residues K321-327); 2) two basic residues of the second Ca^2+^-binging loop (CBL, residues K366 and K369); and 3) two basic residues at the opposite tip (RR, residues R398, R399). The PB and CBL motifs consistently maintained salt bridges with PIP_2_ for both Ca^2+^-free and Ca^2+^-bound forms, while the RR motif only formed dynamic salt bridges with PIP_2_ for the Ca^2+^ free C2B module (Fig. S2, D; Fig. 1 A.3, A.4). For Ca^2+^C2B, strong attraction between Ca^2+^ ions and anionic lipids was observed (Fig. 2S E), and two POPS molecules completed the coordination of chelated Ca^2+^ in the end of the trajectory (Fig. 1 A.4).

**Figure 2.**
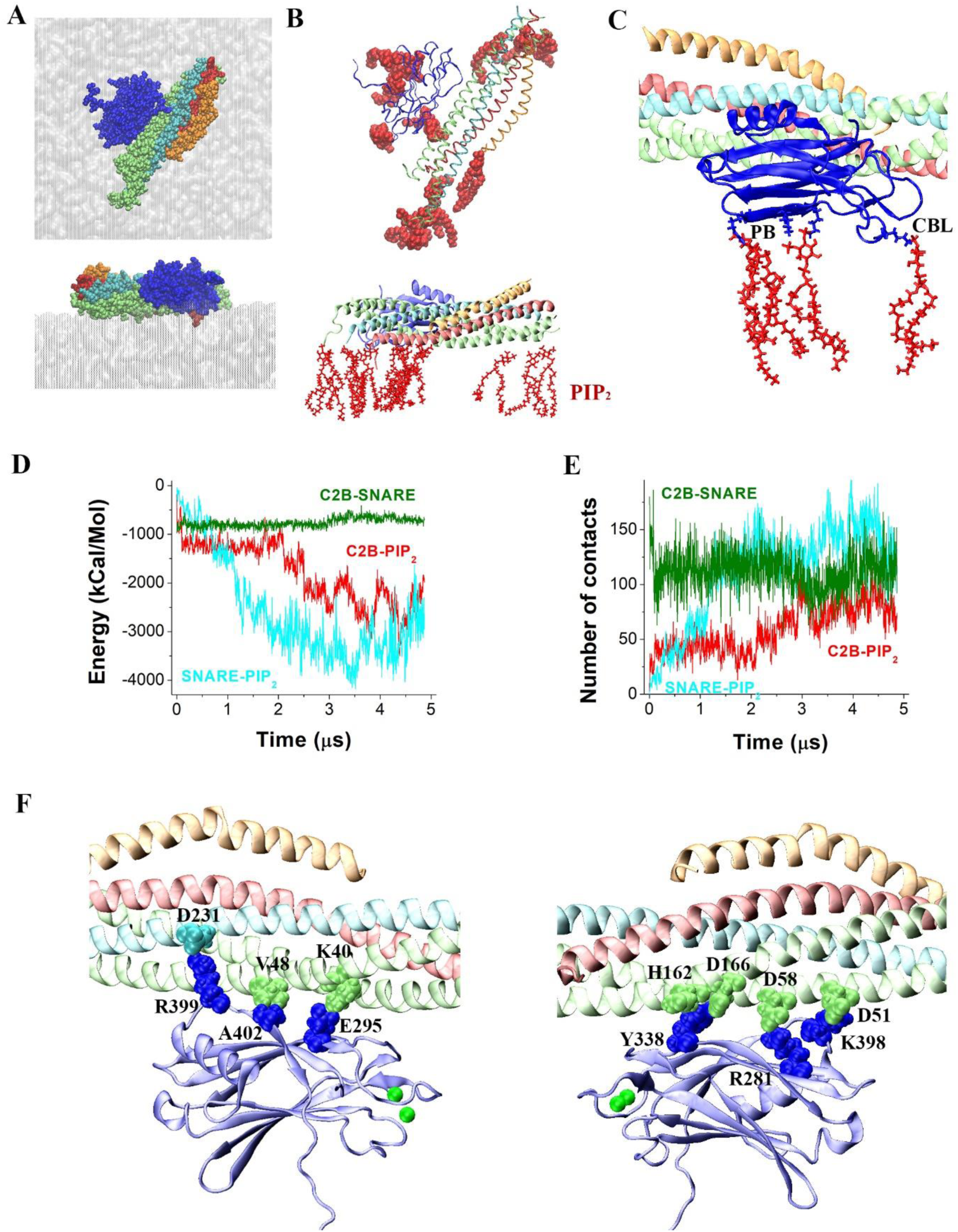
The Ca^2+^C2B-SNARE-Cpx complex anchored at the lipid bilayer imitating PM. **A**. The final point of the MD trajectory: two perpendicular views showing the protein complex in VdW representation on the top of the POPC:POPS:PIP_2_ bilayer (surface representation). Red: Syb; cyan: Syx; lime: SNAP25; orange: Cpx; blue: C2B. Note tight extensive contacts between the protein complex and the bilayer. **B**. The protein complex anchored to multiple PIP_2_ molecules (red). The top view shows PIP_2_ in VdW representation (top) and the side view shows PIP_2_ in bond representation (bottom). **C**. The C2B module is anchored to PIP_2_ via its PB and CBL motifs, as was the caswe for the isolated C2B domain. Ca^2+^ ions are removed for clarity. **D**. The energy of the interactions within the protein-lipid complex along the MD trajectory. Note the energy of PIP_2_-SNARE (cyan) and PIP_2_-C2B (red) decreasing along the trajectory (towards tighter interactions) and then reaching a plateau. Also note a constant energy of C2B-SNARE interactions (dark green), showing that the C2B-SNARE interface largely stayed intact. **E**. The number of VdV contacts within the protein-lipid complex along the MD trajectory. Note numerous contacts between PIP_2_ and SNARE (cyan), PIP_2_ and C2B (red), and C2B and SNARE (dark green) in the end of the trajectory. **F**. The residues forming salt bridges and tight VdW interactions (shown in the VdW representation) between the C2B module and t-SNARE proteins in the end of the trajectory. Two views show the opposite surfaces of the protein complex. Ca^2+^ ions are shown as green spheres.

Thus, both Ca^2+^ free and Ca^2+^ bound forms of the C2B module became strongly attached to the bilayer, although the interactions of the Ca^2+^ bound form were stronger, and evidenced by the shift in energies of the protein-lipid interactions towards more negative values for the Ca^2+^-bound form (Fig. 1 B, C, olive versus purple). However, the attachment configurations differed in the two forms. For the Ca^2+^-free C2B module, the three motifs (CBL, PB, and RR) contributed to dynamic anchoring to the bilayer, producing broad surface attachment (Fig. 1 A.3). In contrast, the Ca^2+^C2B module was tilted, with the Ca^2+^-bound tip penetrating into the bilayer (Fig 1 A.4), Ca^2+^ ions forming coordination bonds with POPS, the CBL motif anchoring to PIP_2_, and the PB region serving as the second anchor.

To evaluate how deeply the protein penetrated into the bilayer, we measured the distance between the lipid surface and the most submerged atom(s) of the protein (Fig. 1 D). Positive values of the penetration parameter correspond to the protein being immersed into lipids, while negative values correspond to the protein being detached from the bilayer. Our results demonstrated that Ca^2+^ binding significantly promotes the lipid penetration, and that the penetration is prominent only for the POPC:POPS:PIP_2_ bilayer imitating PM. As shown in the figure 1D, the Ca^2+^ free form of the C2B module is largely detached from the POPC bilayer, as evident from mostly negative values of the penetration parameter (black line). Ca^2+^ binding brings the C2B domain closer to the POPC bilayer and promotes surface interactions, as illustrated by the penetration parameter fluctuating around zero (green line). The POPC:POPS:PIP_2_ bilayer is consistently penetrated by the Ca^2+^-free C2B domain, with the penetration parameter fluctuating around 1 nm (purple line). Finally, Ca^2+^C2B penetrates the POPC:POPS:PIP_2_ bilayer by approximately 1.5-2 nm (olive line).

Can Ca^2+^C2B penetrate into lipids and simultaneously attach to the SNARE complex? To address this question, we next simulated the interactions of Ca^2+^C2B with the SNARE complex and the lipid bilayer imitating PM. For the initial approximation of the Syt1-SNARE complex, we relied on the crystallography study [27], which identified a large interface between the C2B domain and the SNARE bundle. Since the SNARE complex in its prefusion state is thought to be associated with the fusion affecter Cpx, we generated a model of the Ca^2+^C2B-SNARE-Cpx complex. As decried in the Methods, we combined the structures of Ca^2+^C2B-SNARE [27] with SNARE-Cpx [33, 54] and then positioned the Ca^2+^C2B-SNARE-Cpx above the POPC:POPS:PIP_2_ bilayer (Fig. 3 S A, left) in such a way that the PM and CBL motifs of the C2B module were facing the bilayer.

**Figure 3.**
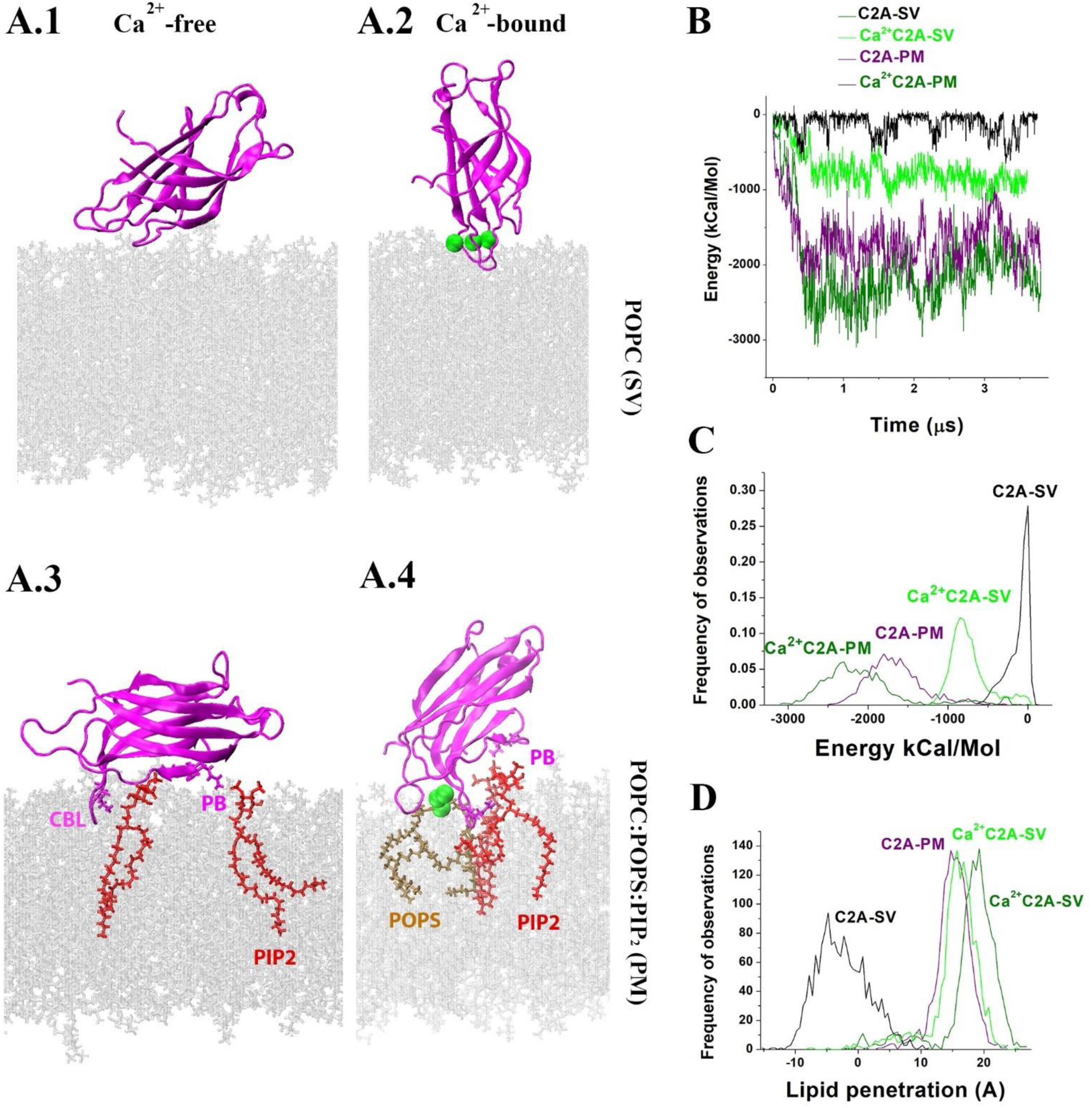
The interactions of the C2A domain with the lipid bilayer. **A**. Representative conformational states in the end of respective MD trajectories showing the attachment of the Ca^2+^ free or Ca^2+^ bound C2A modules to POPC or POPC:POPS:PIP_2_ bilayers. Green spheres depict Ca^2+^ ions. **A**.**1, A**.**2** The C2B module interacts with the SV bilayer via its Ca^2+^-binding tip. **A**.**3**. The C2B module is anchored to PIP_2_ via its PB and CBL motifs. **A**.**4**. CBL of the Ca^2+^C2A module is immersed into the bilayer, being anchored to PIP_2_, while Ca^2+^ ions formed coordination bonds with two POPS molecules. PB is also anchored to PIP_2_. **B**. The energies of the interactions of C2A and Ca^2+^C2A with POPC and POPC:POPS:PIP_2_ bilayers imitating the SV and PM, respectively. The energies are plotted along the respective MD trajectories. **C**. Energy distributions of the interactions of C2A and Ca^2+^C2A with lipid bilayers show that the interactions are cooperatively promoted by Ca^2+^ and PM lipid components (POPS/PIP_2_). **D**. Ca^2+^ and PM lipid components (POPS/PIP_2_) significantly promote the lipid penetration of the C2A module (p<0.01 per K-S test).

The protein complex rapidly attached to the membrane (Fig. 2 A, Fig. S3 A). Over the course of the 4.8 μs MD trajectory, PIP_2_ molecules redistributed towards the protein complex (Fig. S3 B), forming multiple salt bridges with t-SNARE (Fig. 2 B) and the C2B module (Fig. 2 C). The energy of the interactions (Fig. 2 D) and the number of VdW contacts (Fig. 2 E) between PIP_2_ and the proteins plateaued after approximately 3 μs of the simulation and remained at the plateau level until the end of the trajectory. Notably, the tight interface between the C2B domain and the SNARE bundle was not disrupted (Fig. S3 C; Fig. 2 D-F). The salt bridges and tight VdW contacts between the C2B module and the SNARE bundle revealed by crystallography [27] were retained in the end of the trajectory, and in addition new salt bridges between the RR motif of the C2B module and two aspartate residues of t-SNARE (D58 of SNAP 25 and D231 of Syx, Fig. 2F) were formed. These results show that Ca^2+^C2B can simultaneously attach to the SNARE bundle and penetrate into PM.

Thus, both Ca^2+^ free and Ca^2+^ bound forms of the C2B module became strongly attached to the bilayer, although the interactions of the Ca^2+^ bound form were stronger, and evidenced by the shift in energies of the protein-lipid interactions towards more negative values for the Ca^2+^-bound form (Fig. 1 B, C, olive versus purple). However, the attachment configurations differed in the two forms. For the Ca^2+^-free C2B module, the three motifs (CBL, PB, and RR) contributed to dynamic anchoring to the bilayer, producing broad surface attachment (Fig. 1 A.3). In contrast, the Ca^2+^C2B module was tilted, with the Ca^2+^-bound tip penetrating into the bilayer (Fig 1 A.4), Ca^2+^ ions forming coordination bonds with POPS, the CBL motif anchoring to PIP_2_, and the PB region serving as the second anchor.

To evaluate how deeply the protein penetrated into the bilayer, we measured the distance between the lipid surface and the most submerged atom(s) of the protein (Fig. 1 D). Positive values of the penetration parameter correspond to the protein being immersed into lipids, while negative values correspond to the protein being detached from the bilayer. Our results demonstrated that Ca^2+^ binding significantly promotes the lipid penetration, and that the penetration is prominent only for the POPC:POPS:PIP_2_ bilayer imitating PM. As shown in the figure 1D, the Ca^2+^ free form of the C2B module is largely detached from the POPC bilayer, as evident from mostly negative values of the penetration parameter (black line). Ca^2+^ binding brings the C2B domain closer to the POPC bilayer and promotes surface interactions, as illustrated by the penetration parameter fluctuating around zero (green line). The POPC:POPS:PIP_2_ bilayer is consistently penetrated by the Ca^2+^-free C2B domain, with the penetration parameter fluctuating around 1 nm (purple line). Finally, Ca^2+^C2B penetrates the POPC:POPS:PIP_2_ bilayer by approximately 1.5-2 nm (olive line).

Can Ca^2+^C2B penetrate into lipids and simultaneously attach to the SNARE complex? To address this question, we next simulated the interactions of Ca^2+^C2B with the SNARE complex and the lipid bilayer imitating PM. For the initial approximation of the Syt1-SNARE complex, we relied on the crystallography study [27], which identified a large interface between the C2B domain and the SNARE bundle. Since the SNARE complex in its prefusion state is thought to be associated with the fusion affecter Cpx, we generated a model of the Ca^2+^C2B-SNARE-Cpx complex. As decried in the Methods, we combined the structures of Ca^2+^C2B-SNARE [27] with SNARE-Cpx [33, 54] and then positioned the Ca^2+^C2B-SNARE-Cpx above the POPC:POPS:PIP_2_ bilayer (Fig. 3 S A, left) in such a way that the PM and CBL motifs of the C2B module were facing the bilayer.

The protein complex rapidly attached to the membrane (Fig. 2 A, Fig. S3 A). Over the course of the 4.8 μs MD trajectory, PIP_2_ molecules redistributed towards the protein complex (Fig. S3 B), forming multiple salt bridges with t-SNARE (Fig. 2 B) and the C2B module (Fig. 2 C). The energy of the interactions (Fig. 2 D) and the number of VdW contacts (Fig. 2 E) between PIP_2_ and the proteins plateaued after approximately 3 μs of the simulation and remained at the plateau level until the end of the trajectory. Notably, the tight interface between the C2B domain and the SNARE bundle was not disrupted (Fig. S3 C; Fig. 2 D-F). The salt bridges and tight VdW contacts between the C2B module and the SNARE bundle revealed by crystallography [27] were retained in the end of the trajectory, and in addition new salt bridges between the RR motif of the C2B module and two aspartate residues of t-SNARE (D58 of SNAP 25 and D231 of Syx, Fig. 2F) were formed. These results show that Ca^2+^C2B can simultaneously attach to the SNARE bundle and penetrate into PM.

### The interactions of the C2A domain and the C2AB tandem with phospholipid bilayers

Next, we asked how the C2A domain of Syt1 adds to the protein-lipid interactions. First, we simulated the interactions of the isolated C2A domain with the phospholipid bilayers POPC and POPC:POPS:PIP_2_. The initial position of the C2A module relative to the bilayer was in all the cases similar to that for the C2B module (Fig. S1 A). The Ca^2+^-free C2A module showed relatively week but consistent interactions with neutral lipids over the entire course of the 3.8 μs MD trajectory, predominantly attaching to the bilayer via its Ca^2+^ binding tip (Fig. S4, A; Fig. 3 A.1). These interactions were even stronger for the Ca^2+^C2A module, with the Ca^2+^ bound tip being in a contact with the bilayer over the major part of the 3.7 μs trajectory (Fig. S4, B, Fig. 3 A.2, B, C). Interestingly, the analysis of energy components showed a fairly strong attraction of Ca^2+^ ions to the POPC bilayer, and the analysis of Ca^2+^ coordination revealed that oxygen atoms of phosphate groups of two POPC molecules completed the coordination of the chelated Ca^2+^ ions (Fig. S4 C).

Next, simulated the interactions of the C2A domain with the POPC:POPS:PIP_2_ bilayer imitating PM. Both Ca^2+^ free and Ca^2+^ bound forms of the C2A module rapidly attached to the bilayer (Fig. S5 A; Fig. 3 A.3, A4), and remained attached over the entire lengths of respective trajectories. The electrostatic attraction of Ca^2+^ ions to the bilayer (Fig. S5 B) substantially added to the energy of the Ca^2+^C2A interactions with lipids (Fig. 3 B, olive). The Ca^2+^-lipid interactions were largely defined by the POPS lipid component (Fig. S5 B), and two POPS molecules completed Ca^2+^ coordination in the end of the trajectory (Fig. 3 A.4).

The C2A domain has a short polybasic stretch (PB, K189-192, Fig. S5 C), and over the course of the trajectory the PB motif of the C2A domain formed salt bridges with PIP_2_ molecules (Fig. S5, D; Fig. 3 A.3, A.4). The basic residues of the CBL motif of the C2A domain (residues R233 and K236) served as the second anchor attaching to PIP_2_ (Fig. S5 C, D; Fig. 3 A.3, A.4).

The enthalpy analysis (Fig. 3 B, C) demonstrated that the interactions with the POPC:POPS:PIP_2_ bilayer were significantly stronger for both Ca^2+^-free and Ca^2+^-bound forms of the C2A module than their interactions with the POPC bilayer. The lipid penetration (Fig. 3 D) was the deepest for the Ca^2+^C2A module immersed into the POPC:POPS:PIP_2_ bilayer. Thus, our simulations showed that the affinity of the C2A domain to lipid bilayers, as well as the penetration of the C2A module into lipids, were collectively promoted by chelated Ca^2+^ ions, anionic lipids, and PIP_2_ molecules, as was the case for the C2B domain (Fig. 1). For both C2A and C2B modules, chelated Ca^2+^ ions formed coordination bonds with anionic lipids and promoted the immersion of the Ca^2+^ bound tips into the bilayer, while several basic residues anchored to PIP_2_.

However, the simulations also revealed several notable differences in the interactions of the C2A and C2B domains with the bilayers. First, we found that the Ca^2+^ binding loops of the C2A domain had weak affinity to neutral lipids, while in contrast the Ca^2+^ free C2B domain appeared to be highly hydrophilic. The Ca^2+^ bound forms of both domains did form week contacts with neutral lipids, but the binding configurations differed. The Ca^2+^ bound tip of the C2A domain consistently formed contacts with the POPC bilayer, which was not the case for the C2B domain. Instead, Ca^2+^C2B interacted with the POPC bilayer via its α-helix at the opposite tip. Both C2A and C2B domains anchored to PIP_2_ molecules, however the anchors of the C2B domain were more extensive. They included basic residues of CBL, the RR motif at the opposite tip, and an extensive PB region (K321-327). The anchors of the C2A domain were less extensive, since they only included CBL and a relatively short PB stretch (K189-192).

We next asked how these properties of the isolated domains would define the interactions of the C2AB tandem with lipid bilayers. The results obtained for the isolated domains predict that the Ca^2+^-free C2AB tandem would interact with neutral lipids (POPC) via the C2A but not the C2B domain. In contrast Ca^2+^C2AB could interact with the POPC bilayer via both domains. To test this prediction, we simulated the dynamics of the C2AB tandem at the POPC bilayer. The initial structures of the Ca^2+^ free and Ca^2+^ bound forms of the C2AB tandem were taken from our earlier MD simulations of the C2AB tandem in the solution [48]. To minimize the initial influence of the interdomain coupling, for the initial approximation we took a transient state with maximally separated domains (Fig. S6 A), extracted from the MD trajectory obtained for Ca^2+^C2AB.

Only two Ca^2+^ ions were chelated by the C2A domain in the C2AB tandem, as was suggested by earlier experimental [55, 56] and computational [48] studies. The initial state of the Ca^2+^-free C2A tandem was obtained by removing the Ca^2+^ ions.

During the course of the 4.7 µs trajectory of the Ca^2+^-free C2AB tandem, its C2B domain went through multiple configurations, interacting with the membrane only occasionally (Fig. S6 B, C), and these transitions were accompanied by several inter-domain rotations (Fig. S6 D). The Ca^2+^-binding tip of the C2A domain attached to the bilayer within the initial 1 µs of the trajectory and continued interacting with lipids until the end of the simulation (Fig. S6 B; Fig. 4 A, B). These dynamics were consistent with the predictions based on the behavior of the isolated C2 modules. Interestingly, the C2A domain within the tandem penetrated into the bilayer deeper that the isolated C2A module (Fig. 4C, top). This was not the case for the C2B domain, which showed similar penetration within the tandem and in isolation (Fig. 4D, bottom). Possibly, the membrane-detached C2B domain imposes steric constrains on the coupled C2A domain, driving it deeper into the bilayer. This result suggests that the coupling of the C2 domains within the C2AB tandem can affect the immersion of each of the domains into the bilayer.

**Figure 4.**
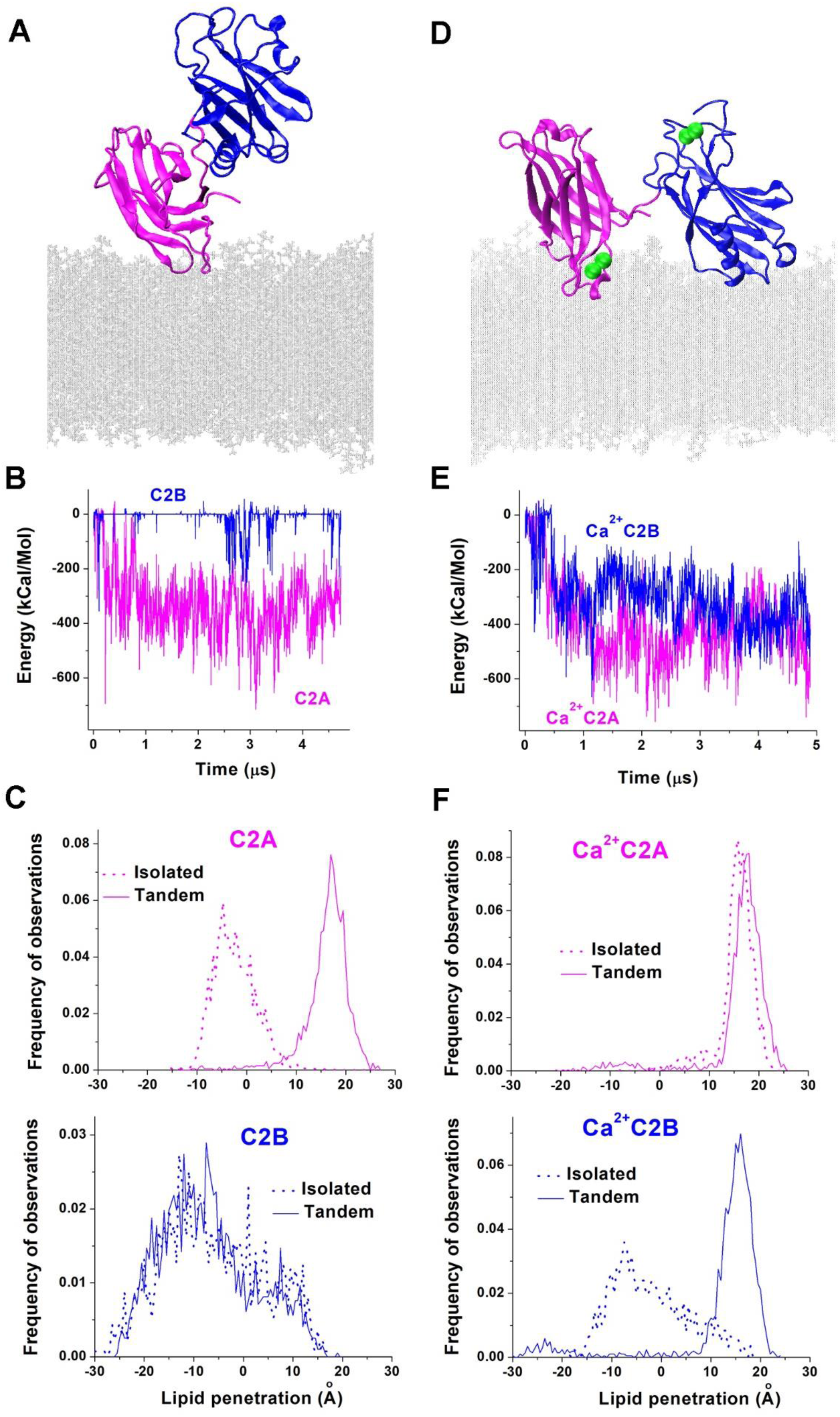
The interactions of the C2AB with neutral lipids (POPC). **A**. The Ca^2+-^free C2AB tandem has its C2A domain (magenta) attached to the bilayer via its Ca^2+-^binding tip, while the C2B domain (blue) is detached from the bilayer. The image shows a representative time-point in the end of the trajectory. **B**. The energy profile of protein-lipid interactions along the trajectory show that the energy of C2B-lipid interactions fluctuates around zero, while the energy of C2A-lipids is consistently negative, showing the attachment of the C2A domain to the bilayer. **C**. The C2A domain penetrates lipids deeper within the C2AB tandem compared to its isolated state (top, p<0.001 per K-S test), while the lipid penetration of the C2B domain is not altered in the tandem (bottom). **D**. Ca^2+^C2AB is attached to the bilayer via the Ca^2+^-bound tip of its C2A domain and the opposite tip of its C2B domain. Green spheres depict Ca^2+^ ions. **E**. Ca^2+^C2A and Ca^2+^C2B have comparable negative energies of the interactions with lipids along the entire trajectory, showing the continuous lipid attachment for both domains. **F**. Both domains penetrate the bilayer deeper being within the tandem, compared to their isolated states, and this effect is more pronounced for the C2B domain. Top: Ca^2+^C2A, p<0.05; bottom: Ca^2+^C2B, p<0.0001.

We next investigated the Ca^2+^C2AB dynamics at the POPC bilayer. After a rapid initial period of instability, the Ca^2+^C2AB tandem adopted a conformation in which Ca^2+^ binding tips of both domains were facing opposite plains (Fig. 4 D, S6 E), and this configuration remained stable for the entire length of the simulation (Fig. 4 E, S6 F, G). The C2A domain interacted with lipids via its Ca^2+^ bound tip, while the C2B domain interacted with the bilayer via its α-helix at the opposite tip (Fig. 4D, S6 E). These results are consistent with the dynamics of the isolated Ca^2+^C2A and Ca^2+^C2B domains on the POPC bilayer, with the isolated Ca^2+^C2A module attaching its Ca^2+^-bound tip to the bilayer (Fig. 3A.2), and the isolated Ca^2+^C2B module attaching its α-helix at the opposite tip (Fig. 1 A.2). This result also agrees with earlier MD simulations, which showed that the Ca^2+^C2AB conformation with Ca^2+^ bound tips facing opposite planes is energetically favorable [48]. Interestingly, C2 domains in a tandem immersed into the bilayer deeper than when being in isolation, showing the synergism, and this was especially pronounced for the C2B domain (Fig. 4 F). These results further illustrate how the coupling of C2 domains could affect their membrane penetration.

We next tested whether this would be the case for PIP_2_-containing anionic lipids. First, we performed MD simulations of the Ca^2+^-free C2AB tandem interacting with the POPC:POPS:PIP_2_ bilayer. The 5.1 μs MD run was started from the same initial state as the previous simulations of the C2AB tandem (Fig. S6 A). During the initial 0.5 μs of the simulation, the tandem underwent two conformational transitions and attached to the bilayer (Fig. S7 A-C). The attached tandem had tightly coupled C2 domains (Fig. 5 A), and each domain was anchored to PIP_2_ in the same way as was observed for the isolated C2 modules. The C2B domain was anchored via its three motifs (PB, CBL, and RR), and the C2A domain was anchored via its two motifs (CBL and PB). The lipid attachments formed by the C2B domain were stronger than those formed by the C2A domain, as evident from more negative energies of the protein-lipid interactions observed for the C2B domain along the trajectory (Fig. 5 B, C). Interestingly, both domains in a tandem formed weaker attachments to the bilayer compared to their isolated states (Fig. 5 C, solid versus dotted lines). Furthermore, the lipid penetration of the C2A domain in a tandem was not as deep as for the isolated C2A domain (Fig. 5 D). In contrast, the lipid penetration of the C2B domain was not affected by the presence of the C2A domain (Fig. 5E). Thus, the tight coupling of the C2 domains in a tandem restricted the interactions of the C2A domain with the bilayer.

**Figure 5.**
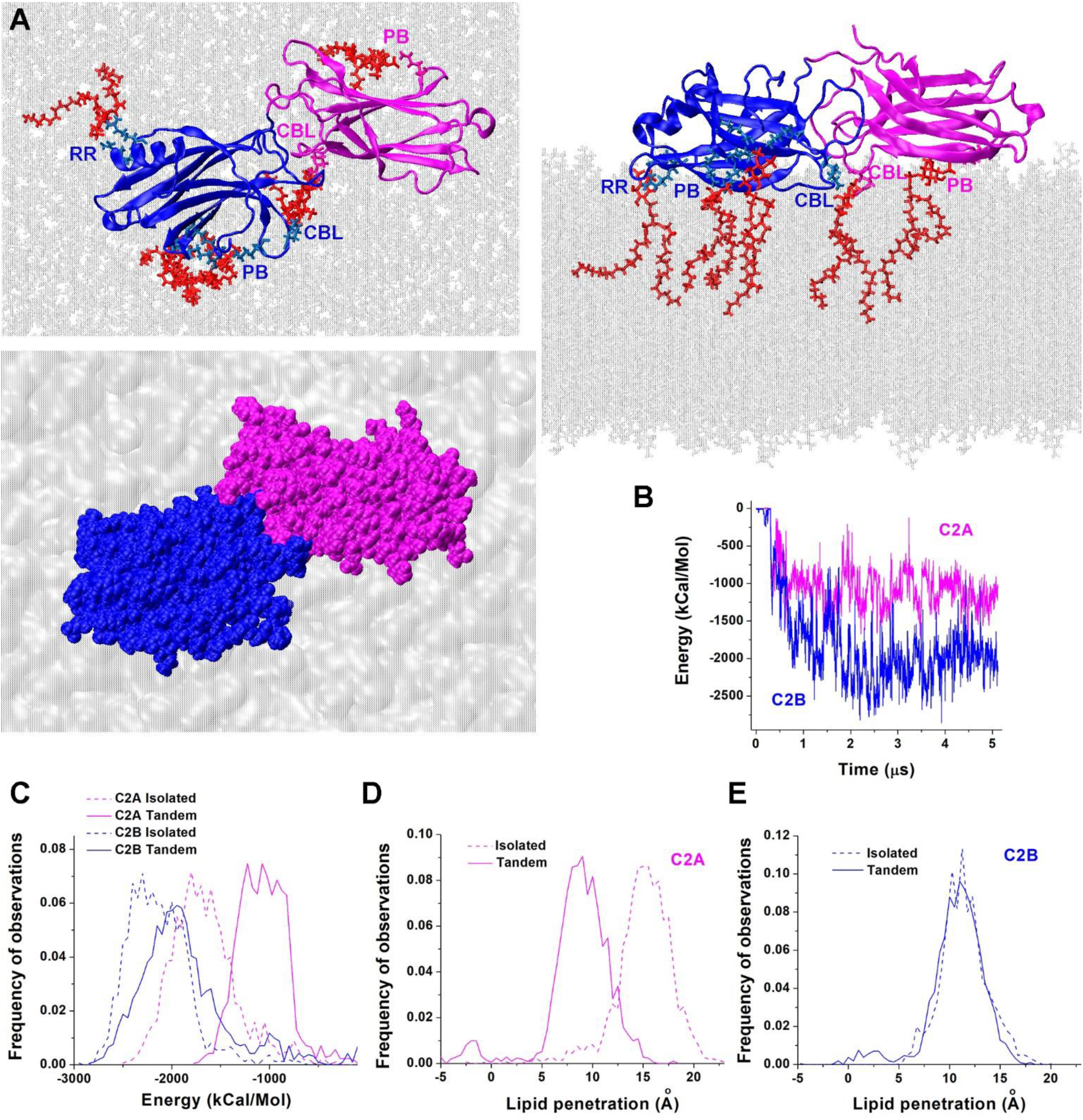
The interactions of the Ca^2+^-free C2AB tandem with the POPC:POPS:PIP_2_ bilayer. **A**. Three views of the protein-lipid complex in the end of the trajectory. Note the PIP_2_ anchor binding simultaneously the CBLs of the C2A and CB domains, as well as additional PIP_2_ anchors (red) binding the PB motifs of the C2 domains and the RR motif of the C2B domain. The VdW representation (bottom) shows a tight coupling between the C2 domains. **B**. Energy profiles of the C2A-lipid and C2B-lipid interactions along the trajectory show lower energies for the C2B domain at the plateau level. **C**. The energies of protein-lipid interactions are shifted towards more positive values for the C2A domain within the tandem, compared to the isolated C2A domain and the C2B domain (p<0.01). **D**. The lipid penetration is significantly reduced for the C2A domain in the tandem compared to the isolated C2A module (p<0.001). **E**. C2B domain penetrates into the lipid bilayer to the same extent, either as a part of the tandem or in isolation.

Next, we investigated the Ca^2+^C2AB tandem attaching to the POPC:POPS:PIP_2_ bilayer. After an initial period of instability of approximately 1.3 μs, the tandem adopted a conformation with perpendicularly oriented domains and maintained this conformation until the end of the 5.1 μs trajectory (Fig. S7 D-F). Both domains attached to the bilayer (Fig. 6 A, Fig. S7 E), and, as was the case for the Ca^2+^ free form, this attachment was stronger for the C2B domain (Fig. 6, B, C). As in the case of the isolated Ca^2+^C2 domains, each domain was attached to PIP_2_ via two anchors: PB and CBL (Fig. 6A). However, the Ca^2+^ ions of both domains remained positioned above the surface of the bilayer (Fig. 6 A, right) and did not form coordination bonds with POPS.

**Figure 6.**
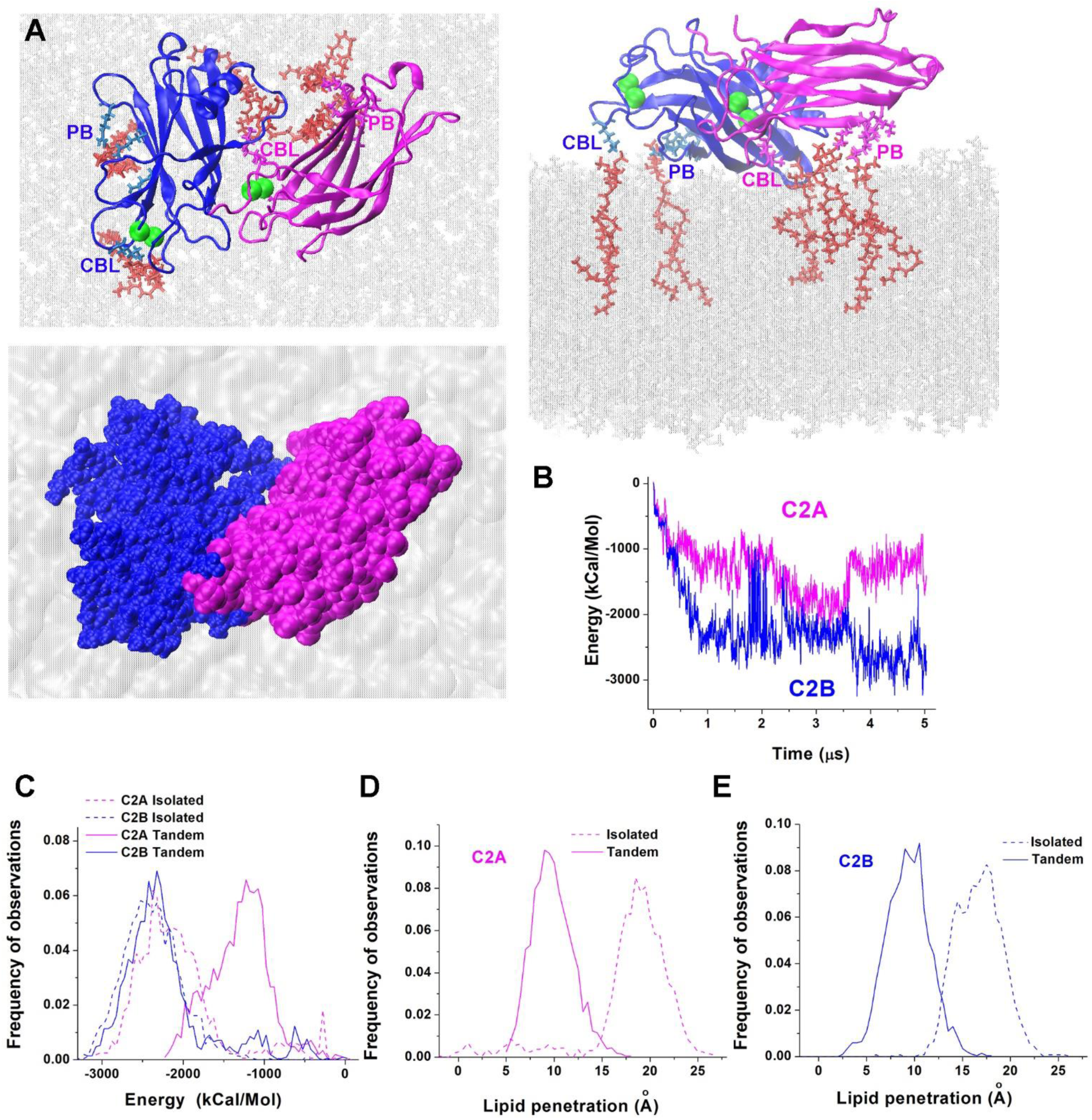
The interactions of Ca^2+^C2AB with the POPC:POPS:PIP_2_ bilayer. **A**. Three views of the protein-lipid complex in the end of the trajectory show PB and CBL anchors attached to PIP_2_ (red). The VdW representation (bottom) shows that C2 domains are tightly coupled. **B**. Energy profiles of Ca^2+^C2A-lipid and Ca^2+^C2B-lipid interactions along the trajectory show more negative energies for the C2B domain. **C**. The energies of protein-lipid interactions are shifted towards more positive values for the C2A domain in the tandem, compared to the isolated C2A domain, as well as to the C2B domain (p<0.001). **D**. The lipid penetration is significantly reduced for the C2A domain in the tandem compared to the isolated C2A module (p<0.001). **E**. The lipid penetration is significantly reduced for the C2B domain in the tandem compared to the isolated C2B module (p<0.01).

Consistently, the energy of the Ca^2+^C2A-lipid interactions was shifted towards more positive values for the tandem, compared to the isolated Ca^2+^C2A module (Fig. 6 C, magenta line). Notably, for both C2 domains within the tandem the lipid penetration was not as deep as for the isolated modules (Fig. 6 D, E).

These results show that the coupling of the C2 domains would counteract the immersion of each domain into the POPC:POPS:PIP_2_ lipid bilayer. This surprising finding suggest that each of the Ca^2+^ bound C2 modules would be more efficient in immersing into PM in isolation, compared to the C2 domains coupled within the C2AB tandem, raising the question of why both domains are critical for triggering fusion. One possibility is that in the prefusion protein-lipid complex the C2 domains of Syt1 become uncoupled. Another possibility is that the immersion of the C2 domains into PM is not the major mechanism by which Syt1 triggers fusion. For example, the coupled C2 domains could trigger fusion by bridging and merging PM and SV via simultaneously penetrating into the opposing bilayers

To investigate the latter possibility, we simulated the dynamics of the C2AB tandem between two bilayers imitating PM and SV. To simulate the two bilayers, we took advantage of the periodic boundary conditions. The advantage of such a system is that the distance between the bilayers is fixed, and the size of the system is kept at a minimum. For the starting configuration, we took the trajectory end-point for Ca^2+^CAB tandem on the POPC bilayer. One of the bilayer leaflets was substituted by the POPC:POPS:PIP_2_ monolayer, and Z-dimension of the cell was adjusted to allow the protein making VdW contacts with the POPC:POPS:PIP_2_ periodic image (Fig. 7 A.1). During the initial 1 μs of the MD run, the Ca^2+^C2B domain attached to the POPC:POPS:PIP_2_ bilayer (Fig. 7 A.2), being anchored to a PIP_2_ molecule via its CBL and PB motifs. The contacts between Ca^2+^C2A and POPC remained extensive (Fig. 7 A.2, B), and these interactions created a prominent protrusion in the POPC lipid bilayer at the interface of C2 domains (Fig. 7 A.2, arrow). Hypothetically, such protrusions could promote the stalk formation and lipid merging. However, subsequent simulations produced a conformational transition of the Ca^2+^C2A domain, followed by its attachment to the POPC:POPS:PIP_2_ bilayer (Fig. 7 A.3, B) via its CBL and PB motifs. Meanwhile, additional PIP_2_ molecules aggregated and attached to the Ca^2+^C2B domain (Fig. 7 A.3). This configuration proved to be stable for the subsequent 5.5 us of the simulation, suggesting that the PM-SV bridging by the Ca^2+^C2AB tandem is a transient state, and that both domains would tend to interact with PM.

**Figure 7.**
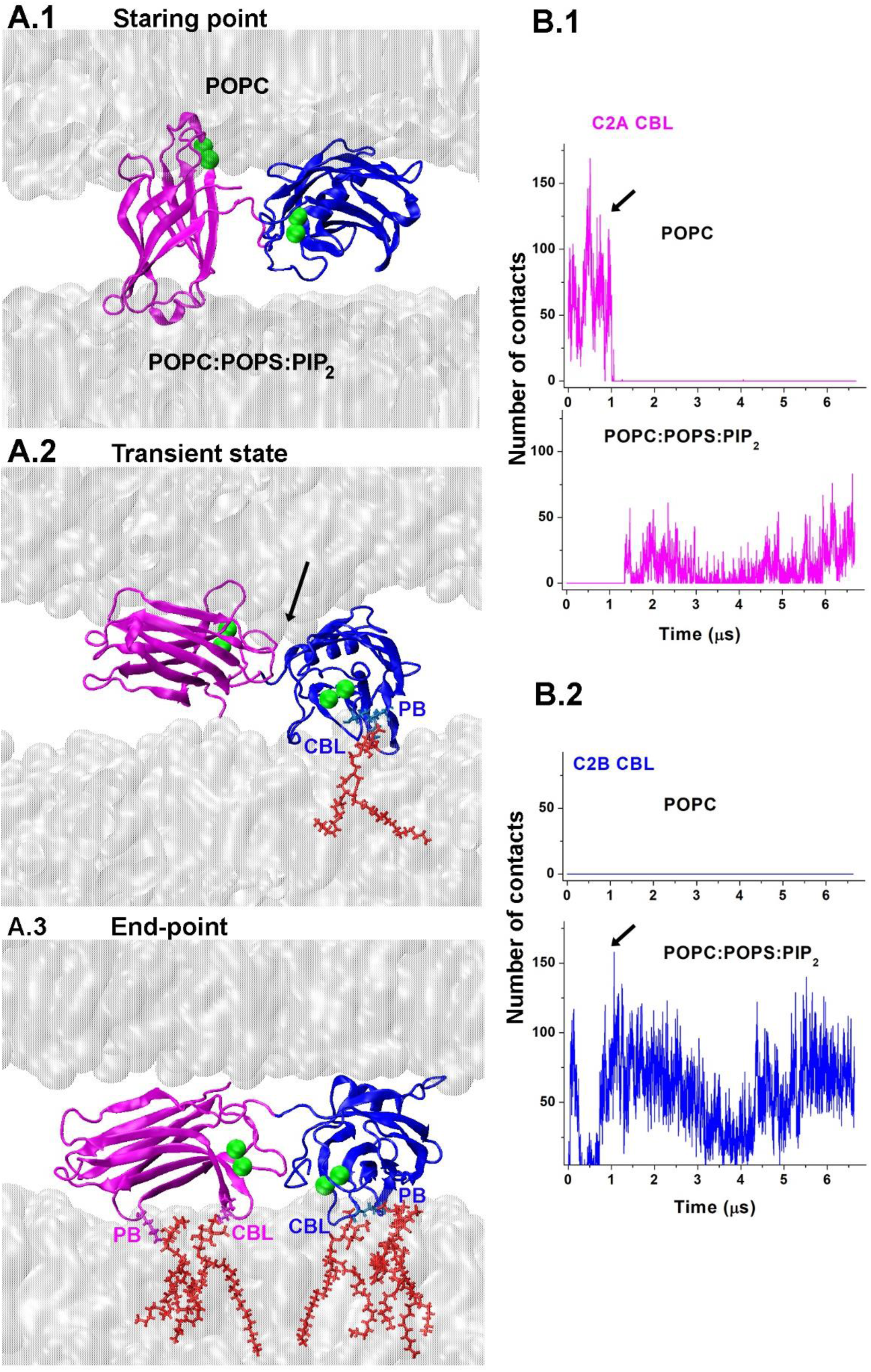
The dynamics of the Ca^2+^C2AB tandem between the lipid bilayers imitating PM and SV. **A**. Three conformations along the trajectory. **A**.**1**. The initial configuration. The CBL motif of Ca^2+^C2A is immersed into the POPC bilayer. A.2. The transient state (1 μs). The Ca^2+^C2B is anchored to PIP_2_ (red) via its CBL and PB motifs. This state is characterized by a prominent lipid protrusion (arrow). **A**.**3**. The final state. Both C2 domains are anchored to PIP_2_ clusters via their CBL and PB motifs. **B**. The number of contacts between lipids and Ca^2+^-binding loops of each C2 domain along the trajectory (residues 170-180 and 230-240 for the C2A domain; residues 300-310 and 360-370 for the C2B domain). Note the conformational transition after the initial 1 μs. **B**.**1**. The Ca^2+^-binding loops of the C2A domain form numerous VdW contacts with the POPC bilayer during the initial 1 μs of the trajectory. Subsequently, the contacts are formed only with the opposing POPC:POPS:PIP_2_ bilayer. **B**.**2**. The Ca^2+^-binding loops of the C2B domain form VdW contacts only with the POPC:POPS:PIP_2_ bilayer. Arrow in the panels B.1 and B.2 points to the time-point shown in A.2.

Thus, our simulations revealed a transient state of the Ca^2+^C2AB tandem, which could bridge the SV and PM bilayers and promote lipid protrusions potentially leading to a stalk formation. However, the simulations also showed that this state would compete with a more favorable and stable state in which both domains would attach to PM via their CBL and PB motifs. The question then still remains: how do the C2 domains cooperate in promoting fusion? To elucidate this issue, we investigated whether the attachment to the SNARE complex could alter the dynamics of the C2AB tandem and its interactions with PM.

### Interactions of the C2AB tandem with the SNARE-Cpx complex and phospholipids

We started from the structure of the C2AB-SNARE complex obtained by crystallography [27]. This complex has an extensive C2B-SNARE interface (I1), and our simulations of the C2B-SNARE-Cpx complex (Fig.2) demonstrated that this interface is stable in the water/ion/lipid environment. To understand how the C2A domain adds to the protein-lipid interactions, we constructed the C2AB-SNARE-Cpx complex by combining C2AB-SNARE [37] with SNARE-Cpx [54] as described in Methods. The structure was then equilibrated, and the 9 µs MD run was performed (Fig. S8). Surprisingly, we observed a sharp increase in the energy of C2B-SNARE interactions during the initial 500 ns of the trajectory (Fig. S8 A), denoting weakened SNARE-C2B interactions. Notably, such increase was not observed in the absence of the C2A domain (Fig. 2D, olive). Furthermore, the energy distribution for C2B-SNARE interactions was shifted towards more positive values in the presence of the C2A domain (Fig. S8 B), suggesting that the interactions of the C2A domain destabilized the complex. The analysis of the trajectory showed that the complex underwent a conformational transition between 4.8 and 5.0 µs (Fig. S8 C), so that the C2B module shifted along the SNARE bundle towards its membrane-distal terminus, forming a new interface with t-SNARE proteins (I2, Fig. S8 C). This new conformation corresponded to a local minimum in the energy of the C2B-SNARE interactions (Fig. S8 A, I2). Together with the results presented in figure 2, these results indicate that although the C2B-SNARE interface I1 [37] is energetically favorable and likely stable in the water/ion environment, the position of the C2A domain obtained by crystallography may not be energetically optimal at physiological ion concentrations. Furthermore, optical studies showed that the Syt1-SNARE complex samples multiple conformational states in the solution [57], and NMR approach identified the structure of the Syt1-SNARE complex in which the PB motif of the C2B domain anchored onto the SNARE bundle [28]. We therefore sought to employ MD simulations to investigate the conformational states of the C2AB-SNARE-Cpx complex more systematically.

We generated four initial approximations for the C2B-SNARE interface (Fig. S9): I1 interface obtained by crystallography [27]; I2 interface produced by our initial MD simulations (Fig. S9); I3 interface obtained by NMR studies [28]; and I4 interface obtained by docking of the C2B domain to the SNARE bundle basing on spectroscopy [58] and genetic [47] studies, as described in Methods. All the complexes had the conformation of the C2AB tandem with perpendicularly oriented C2 domains (Fig. S10 A), since both crystallography [55] and MD [48] studies suggest that this conformation would be energetically favorable. Each complex was equilibrated, and the production MD runs were performed. In all the four trajectories (Fig. S10 B), both C2 modules were attached to the SNARE-Cpx bundle (note negative energies for each of C2 domains interacting with the SNARE-Cpx bundle), however, the interactions of the C2B domain prevailed (note lower energies for C2B). In each run, we ensured that the energies of the interactions of the SNARE-Cpx bundle with C2 domains reached a plateau (of at least 2 μs), and this required MD runs of 6-9 μs each (Fig. S10 B, I1-I3). The only exception was the state I4, which converged to the state similar to the final point of the I2 trajectory (I2^f^, Fig. S10 C). Respectively, the I4 run was terminated (at 2.7 μs) as being redundant. Thus, four initial approximations of the C2AB-SNARE-Cpx complex, built upon four different C2B-SNARE interfaces (Fig. S9, S10A) converged into three different conformational states (I1^f^, I2^f^, I3^f^, Fig. S10 C; Fig. 8 A), which were stable at a microsecond scale.

**Figure 8.**
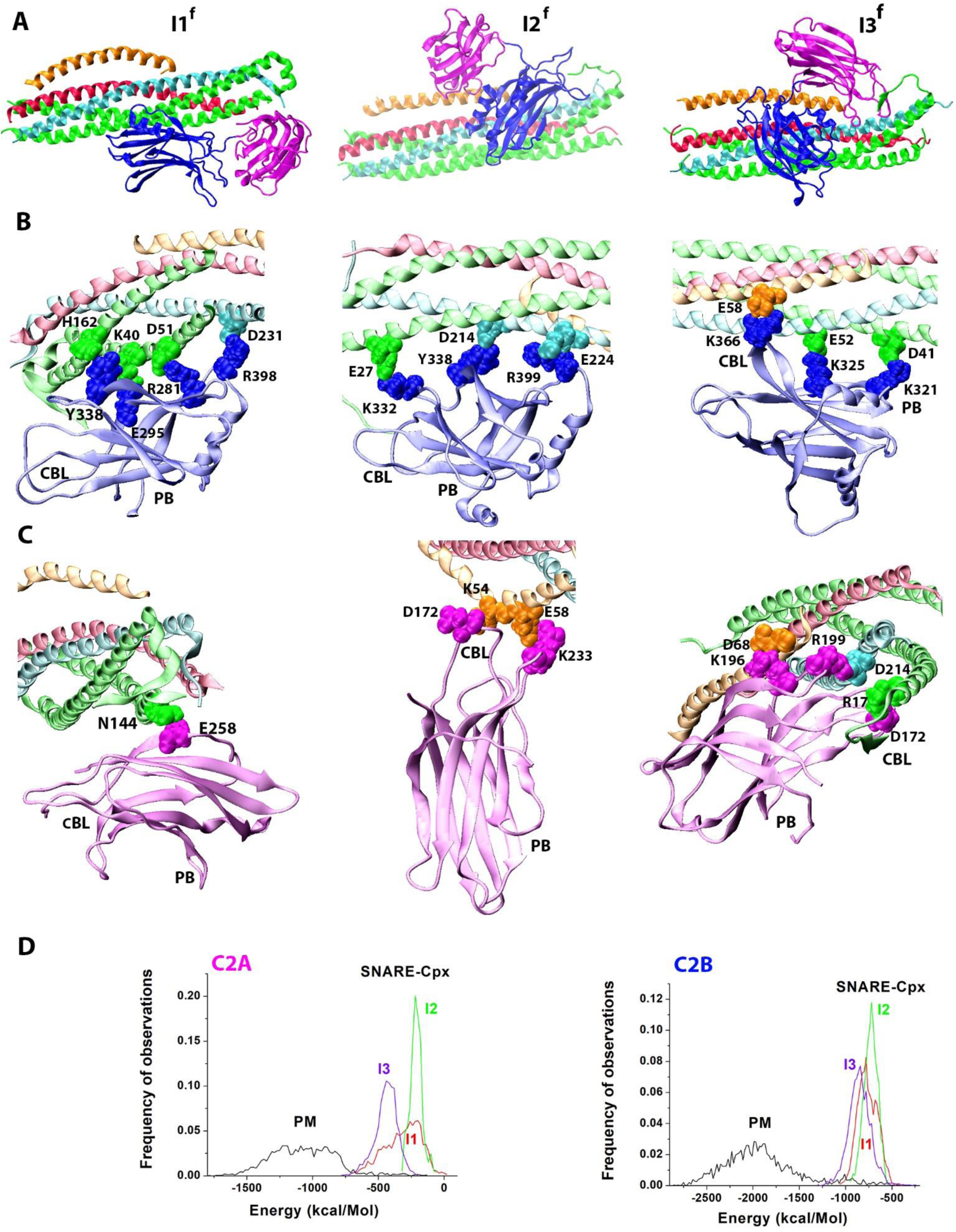
Three conformational states of the C2AB-SNARE-Cpx. **A**. The overall topology of the conformational states I1^f^, I2^f^, and I3^f^. Red: Syb; cyan: Syx; green: SNAP25; orange: Cpx; blue: C2B; magenta: C2A. **B**. The interface between the C2B domain and the SNARE-Cpx bundle for the three states. The residues forming salt bridges and the strongest VdW attachments are shown in the VdW representation and marked. **C**. The interface between the C2A domain and the SNARE-Cpx bundle. The locations of the CBL and PB motifs are marked for both C2 domains. **D**. Both C2 domains have higher afffinities to PM than to the SNARE-Cpx bundle. The graphs depict the energy distributions for the interations of the C2A or C2B domain with the SNARE-Cpx bundle derived for each of the three complexes over the final 2 μs of the respective trajectories. For the comparison, the energy distributions for the interations of the C2 domains with the lipid bilayer imitating PM (POPC:POPS:PIP_2_) are shown (black lines, re-plotted from Fig. 5 C). Note that the PM distributions are shifted towards more negative energy values and largely do not overlap with the SNARE-Cpx distributions, showing a strong preference for each of the C2 domains to attach to PM versus the SNARE-Cpx bundle.

The I1 interface between the C2B domain and the SNARE bundle [27] remained largely unchanged over the course of the trajectory (I1, Fig. S10 A versus C, note the unchanged position of the C2B domain). The final endpoint complex (I1^f^, Fig. 8 A) had several stabilizing salt bridges between the C2B domain and t-SNARE (Fig. 8 B: R281 of C2B and D51 of SNAP25; E295 of C2B and K40 of SNAP25; R398 of C2B and D231 of Syx), as well as multiple VdW contacts (note Y338 of C2B and H162 of SNAP25, Fig. 8 B, I1^f^). Notably, mutating the C2B residues R281, E295, and Y338 belonging to this interface produces Syt1 loss-of-function [47]. The interactions of the C2A domain were weaker and involved only single salt bridge with SNAP25 in the end of the trajectory (Fig. 8 C, I1^f^).

In the I2^f^ complex, the C2B module was shifted towards the membrane-distal end of the SNARE bundle, compared to the I2^f^ complex. Interestingly, the C2A module underwent a conformational transition in the beginning of the I2 trajectory (at 1.7 μs) and formed two salt bridges with Cpx, which were stable until the end of the trajectory (for over 5 μs). The salt bridges were formed between Ca^2+^-binding loops of the C2A domain and the Cpx stretch connecting it central and accessory helixes (Fig. 8 C, I2^f^). Thus, the MD simulations revealed a new tri-partied interface between Syt1, Cpx, and the SNARE bundle, in which the C2B domain forms a tight interface with t-SNARE, while the C2A domain forms salt bridges with Cpx (Fig. 8 A-C, I2^f^).

The I3 complex had the PB stretch of the C2B domain binding t-SNARE [28], and these interactions stabilized over the curse of the 6 µs MD trajectory. In addition, the CBL motif of the C2B domain formed a salt bridge with Cpx (Fig. 8 B, I3^f^). The C2A domain although formed a salt bridge with Cpx via the 3^rd^ loop of its Ca^2+^-binding tip (K196 of C2A and D68 of Cpx, Fig. 8 C, I3^f^). This loop of the C2A domain also formed a salt bridge with Syx (R199 of C2A and D214 of Syx), while the first Ca^2+^-binding loop of the C2A domain formed a salt bridge with SNAP25 (D172 of C2A and R17 of SNAP25). Thus, both C2 domains in the I3^f^ complex formed tight links with the SNARE-Cpx bundle, and these interactions involved the CBL and PB motifs of the C2B domain, and well as the Ca^2+^-binding tip of the C2A domain.

The enthalpy analysis revealed that the interactions between Syt1 and the SNARE-Cpx bundle are most favorable energetically for the complex I3^f^. The energy of the interactions between each of the C2 domains and the SNARE-Cpx bundle was shifted towards more negative values for the complex I3^f^ (Fig. 8D), suggesting that the I3^f^ complex is likely to be prevalent in the intracellular saline. This finding agrees with the results of the NMR studies [28]. However, the proximity to PM may alter the interplay between the conformational states of the Syt1-SNARE-Cpx complex. Indeed, the comparison with C2-PM interactions (Fig. 8 D, black lines) shows that the attraction to PM is much stronger for each of the C2 domains than the attraction to the SNARE-Cpx bundle. Therefore, in the proximity to PM, the conformational state(s) of the Syt1-SNARE-Cpx complex enabling simultaneous interactions of C2 domains with PM would prevail.

Out of the three conformational states (Fig. 8) identified by our simulations, the state I1^f^ is the most likely candidate to satisfy this requirement. Indeed, the formation of the I3^f^ complex involves the CBL and PB motifs of the C2B domain, which are also required for the attachment of the C2B domain to PM. Therefore the formation of the I3^f^ complex would preclude the attachment of the C2B domain to PM. The formation of the I2^f^ complex does not involve either CBL or PB of the C2B domain, however it involves CBL of the C2A domain. In contrast, the I1^f^ complex does not involve either CBL or PB motifs of either of the C2 domains, suggesting that the I1^f^ conformational state of the C2AB-SNARE-Cpx complex would allow Syt1 to simultaneously attach to lipid bilayers.

To test this suggestion, we positioned the I1^f^ complex over the lipid bilayer imitating PM (POPC:POPS:PIP_2_), substituted the Ca^2+^ free form of the C2AB tandem by its Ca^2+^-bound form, equilibrated this structure in water/ion environment, and performed the 8.4 µs MD run. Over the course of the trajectory, both Ca^2+^ bound tips immersed into the bilayer (Fig. 9 A, B). Notably, the I1 interface between the C2B domain and the SNARE bundle was not disrupted or even weekend. C2B domain remained tightly linked to the bundle via several salt bridges and hydrophobic interactions with t-SNARE proteins (Fig. 9 C). In contrast, the C2A domain separated from the bundle in the course of the simulation (Fig. 9 A, B). Notably, PIP_2_ molecules clustered around the protein complex over the course of the trajectory, anchoring both C2 domains and the SNARE bundle (Fig. 9 D). These results show that the interactions of the C2 domains with the bilayer do not compromise the ability of the C2B domain (but not the C2A domain) to form tight interactions with the SNARE bundle via the I1 interface.

**Figure 9.**
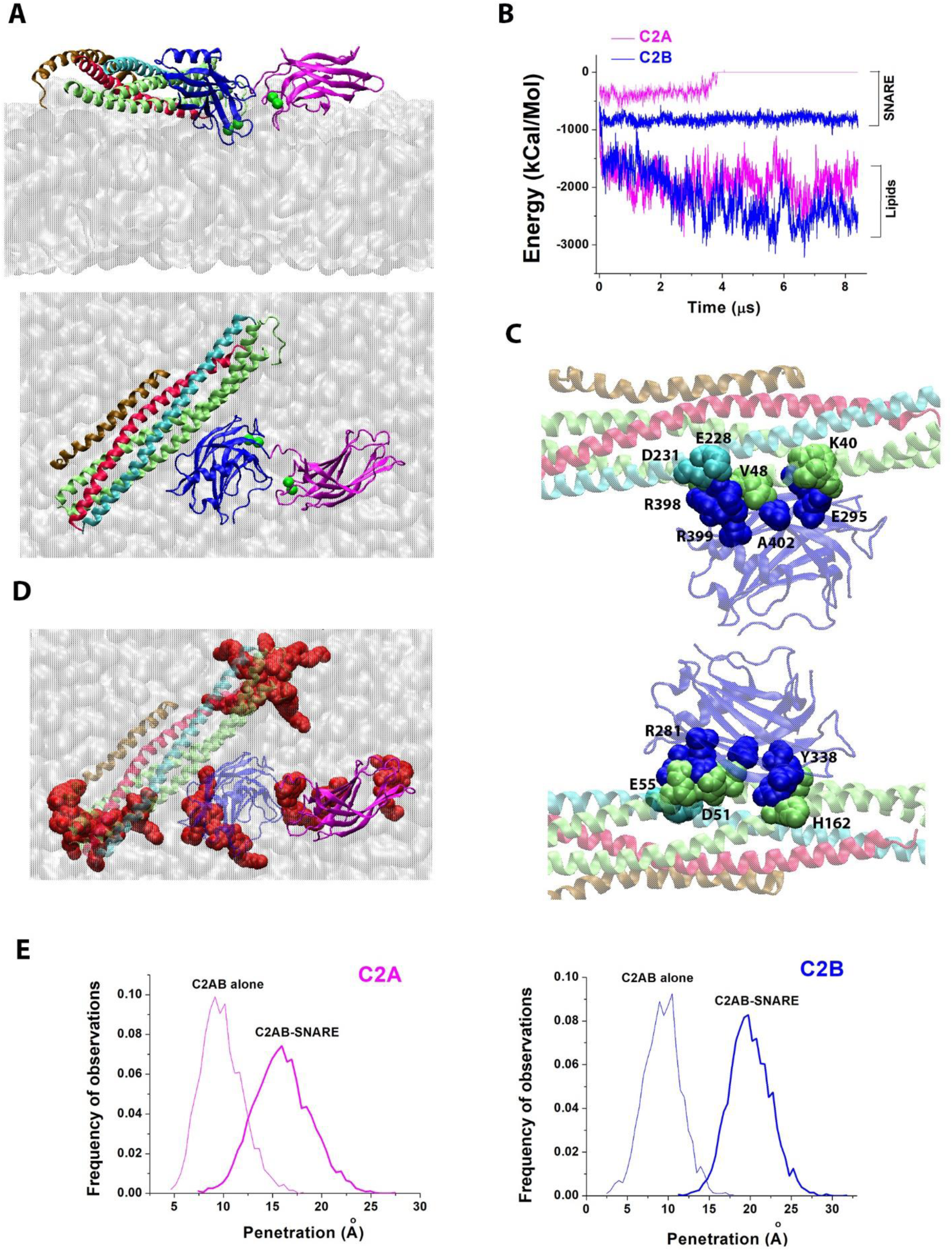
The association of the Ca^2+^C2AB tandem with the SNARE-Cpx bundle promotes the immersion of the Ca^2+^-bound tips of C2 domains into the lipid bilayer imitating PM. **A**. Two perpendicular views showing the Ca^2+^I1-PM complex at the trajectory end-point. Note that the Ca^2+^ -bound loops of both domains are immersed into the bilayer (top), and that the C2B domain is attached to t-SNARE (bottom). C2A - magenta; C2B - blue; Ca^2+^-green spheres; Syb - red; Syx - cyan; SNAP25 - lime; Cpx – ochre; POPC:POPS:PIP_2_ bilayer - silver, surface representation. **B**. The energy profile along the trajectory for the interactions of each of the C2 domains with the SNARE bundle and with the lipid bilayer. Note the steady level the C2B-SNARE attractions. In contrast, the energy of the C2A-SNARE interactions increases to zero at the 4 μs time-point, denoting the separation of the C2A domain from the SNARE-Cpx bundle. Also, note the decrease in the energies of C2-lipids interactions, denoting the immersion of C2 tips into the bilayer. **C**. Two opposite views showing multiple salt bridges and tight VdW contacts between the C2B domain and the SNARE bundle. Ca^2+^ ions are removed for clarity. **D**. PIP_2_ molecules (red, VdW representation) are clustered around the protein complex, anchoring the C2 domains and t-SNARE. **E**. The Ca^2+^-bound tips of the C2 domains penetrate the bilayers to a significantly larger extent than in the absence of the SNARE bundle (p<0.01 for C2B, p<0.001 for C2B). The graphs show the frequency distributions for the lipid penetration. For comparison (thin lines) we re-plotted the distributions derived for the Ca^2+^C2AB tandem alone interacting with the POPC:POPS:PIP_2_ bilayer (the same data as in Fig. 6 D for the tandem).

We next assessed how the C2 domains penetrate into lipids within this protein-lipid complex (Ca^2+^I1-PM). Strikingly, we found that the lipid penetration for each of the C2 domains within the Ca^2+^I1-PM complex was significantly deeper than the lipid penetration for the C2 domains within the isolated C2AB tandem interacting with PM (Fig. 9 E). In other words, the attachment of the C2B domain to the SNARE bundle promoted the immersion of both C2 domains into the lipid bilayer. The likely explanation for this result is that the tight coupling between the C2B domain and the SNARE bundle within the Ca^2+^I1-PM complex uncouples the C2A and C2B domains within the C2AB tandem and allows them to deeper penetrate into lipids.

Thus, our results show that the interactions with the SNARE bundle promote the conformation of the C2AB tandem with uncoupled C2 domains, which is optimal for the lipid penetration. In the absence of the SNARE bundle, the C2 domains within the C2AB tandem mutually hinter their penetration into lipids (Fig. 6). However, the attachment of the C2B domain to the SNARE bundle via the I1 interface uncouples the C2 domains and abolishes their mutual hindering, thus enabling the Ca^2+^-bound tips of the C2 domains to penetrate into PM to the extent of the isolated C2 domains. The distortion, curvature, and tension produced by the immersion of C2 domains into PM could drive fusion synergistically with SNARE zippering.

### The model of the pre-fusion protein-lipid complex

We next investigated whether the Ca^2+^I1-PM complex could drive the SV-PM fusion. To test this, we added to the Ca^2+^I1-PM complex transmembrane domains of Syb and Syx embedded in the lipid bilayer imitating SV. To generate this molecular system, we performed the following steps (Fig. S 11). First, we took the X-ray structure [59] of the SNARE complex with intact transmembrane domains of Syb and Syx (3IPD, Fig. S11 A) and removed SNAP25. Next, we generated the Syx^tm^-Syb^tm^ complex (Fig. S11 B) with bent transmembrane domains. This was done by keeping all the atoms of the coil-coiled region fixed, imposing distance constraints on C-terminus Cα atoms of Syx^tm^ and Syb^tm^, and performing the Monte-Carlo/minimization procedure with ZMM software [60]. Next, the structures of Syx and Syb within the Ca^2+^I1-PM complex (Fig. 9) were sequentially replaced by Syx^tm^ and Syb^tm^, respectively, employing the RMSD alignment. The resulting system (Fig. S11 C) had Syx^tm^ partially embedded in the lipid bilayer imitating PM. The lipid monomers overlapping with the Syx^tm^ were removed from the system. Finally, the second membrane patch (POPC) imitating an SV was added to the system and positioned parallel to the POPC:POPS:PIP_2_ bilayer at a distance allowing VdW contacts with the C2AB tandem (Fig. S13 D). The lipid monomers of the POPC patch overlapping with Syb^tm^ were then removed from the system. Water and ions were added, and Z-dimension of the periodic cell was adjusted to allow a gap of 4.5 nm between the POPC patch and the periodic image of the POPC:POPS:PIP_2_ patch (Fig. S11 E, left). The size of the POPC patch was adjusted in such a way that its Y dimension aligned with the POPC:POPS:PIP_2_ patch, while its X dimension was shorter by 1.5 nm (Fig. S13 E, right). This was done to allow water molecules to circulate freely, so that artificial pressures between the bilayers are not generated.

Following equilibration of this system, we performed a 6 μs production MD run. Notably, in the course of the simulations we observed multiple instances of the two bilayers approaching each other and making extensive contacts (Fig.10 A, B). Merging the membranes was usually associated with noticeable curvature of the bilayer imitating PM (Fig. 10 A, right, arrow). We also observed numerous transient zippering events for the Layer 9 of the SNARE complex (Fig. 10 C), which was initially unzippered. Over the entire course of the trajectory, the Ca^2+^-bound tips of both C2 domains were immersed into the PM bilayer (Fig. 10 D), with a deeper penetration for the C2B domain. The C2B domain also remained attached attached to the SNARE bundle (Fig. 10 E). These results illustrate how the immersion of the C2 domains of Syt1 into PM synergistically with SNARE zippering could bring together PM and SV and promote their merging.

**Figure 10.**
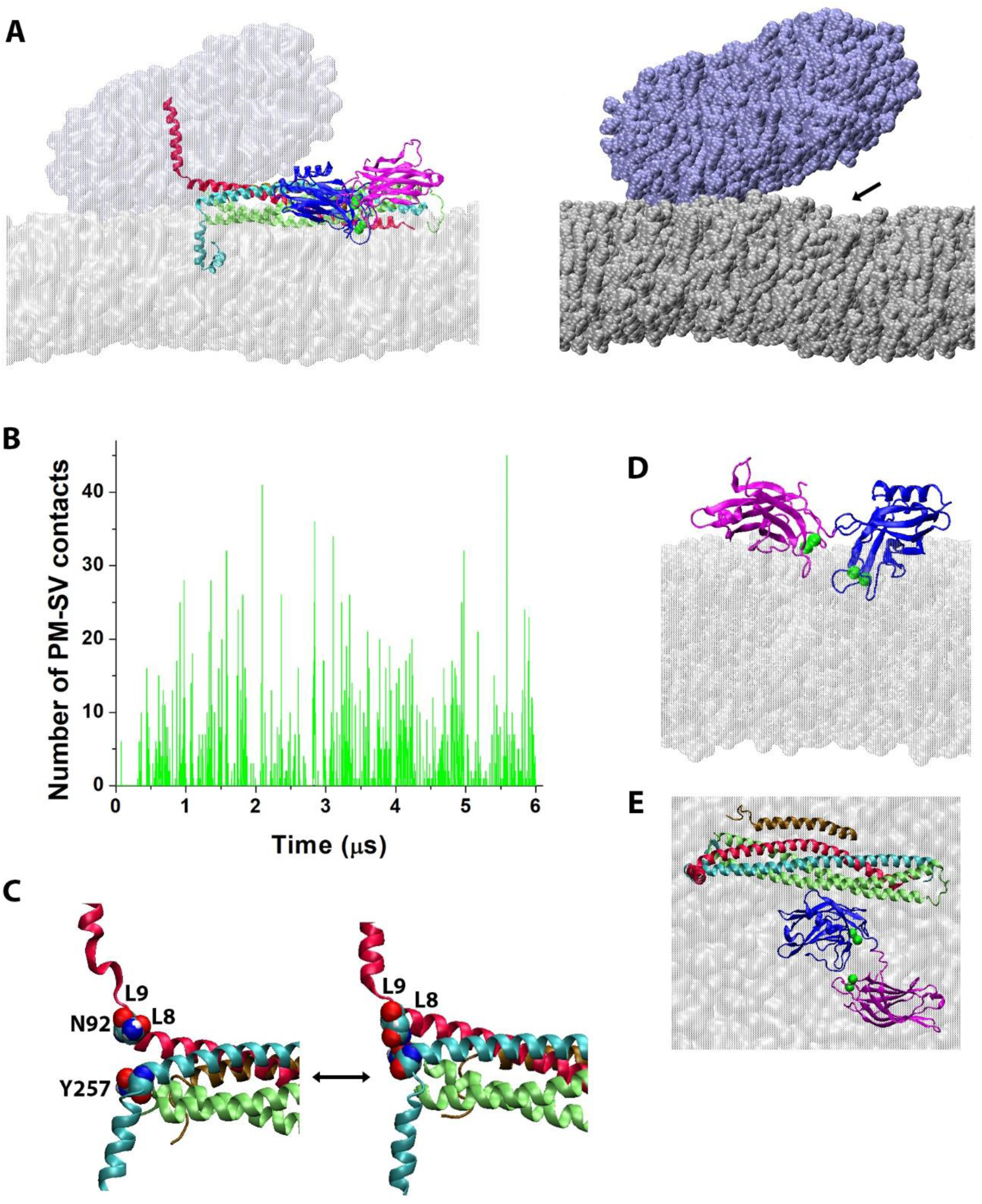
The prefusion Ca^2+^Syt1-SNARE complex promotes lipid merging. **A**. Two views of the protein-lipid complex Ca^2+^I1-PM-SV in the end of the trajectory. The membranes imitating PM (silver) and SV (ice-blue) form extensive VdW contacts. Left: The Syb (red) transmembrane domain spans through SV, and the Syx (cyan) transmembrane domain spans through PM, while the Ca^2+^-bound C2A (magenta) and C2B (blue) domains of Syt1 immerse into PM. Two membranes (surface representation) form extensive contacts. Right: The protein complex is removed for clarity and the membranes are depicted in VdW representation to show numerous VdW contacts between SV and PM. Note a prominent PM curvature (arrow) at the site of Syt1 binding. **B**. Numerous VdW contacts between the membrane patches imitating PM and SV are repeatedly formed over the course of the trajectory. The points of zero contact denote the transient separations of the membrane patches. Note that the overall number of contacts increases over the initial 1 μs of the trajectory, and then the system reaches an equilibrium, with the two membranes repeatedly merging and separating. **C**. Zippering of the Layer 9 of the SNARE complex. The unzippered Layer 9 (left) has the residues N92 of Syb and Y257 of Syx (shown in VdW representation) being separated, while the complex with the fully zippered Layer 9 has these residues forming a hydrogen bond. **D**. The C2AB tandem in the end of the trajectory being immersed into PM. **E**. Top view of the Ca^2+^C2AB-SNARE-Cpx complex. The membrane patch imitating SV is removed for clarity.

To delineate the specific role of Ca^2+^ chelation by Syt1 in this *in silico* system, we repeated the above simulations for the Ca^2+^ -free form of the C2AB tandem. We started from the same initial approximation (Fig. S11) but removed chelated Ca^2+^ ions. The system was subsequently neutralized and re-equilibrated, and the 6 µs production MD run was performed. We observed that over the course of the trajectory the edges of the two membrane patches periodically approached each other, however they usually did not form VdW contacts (Fig. S12 A). Over the entire course of the simulation, we observed only several frames in which VdW contacts were present, and these contacts were not extensive (Fig. S12 B).

The overall number of contacts was drastically diminished for the Ca^2+^ free form of the C2AB tandem, compared to its Ca^2+^ bound form (Fig. 11 A). Importantly, the system with the Ca^2+^-bound form of the C2AB tandem consistently showed relatively long-lasting (for tens of nanoseconds) stretches of the trajectory with the two patches being in contact (Fig. 11 B, green). In contrast, in the absence of Ca^2+^ these contacts were very brief (one or two trajectory frames, corresponding to 0.2-0.5 nanoseconds) and not extensive (Fig. 11 B, violet). Thus, the ability of the Ca^2+^C2AB-SNARE complex to drive lipid merging in our molecular system is not reproduced for the Ca^2+^-free complex.

**Figure 11.**
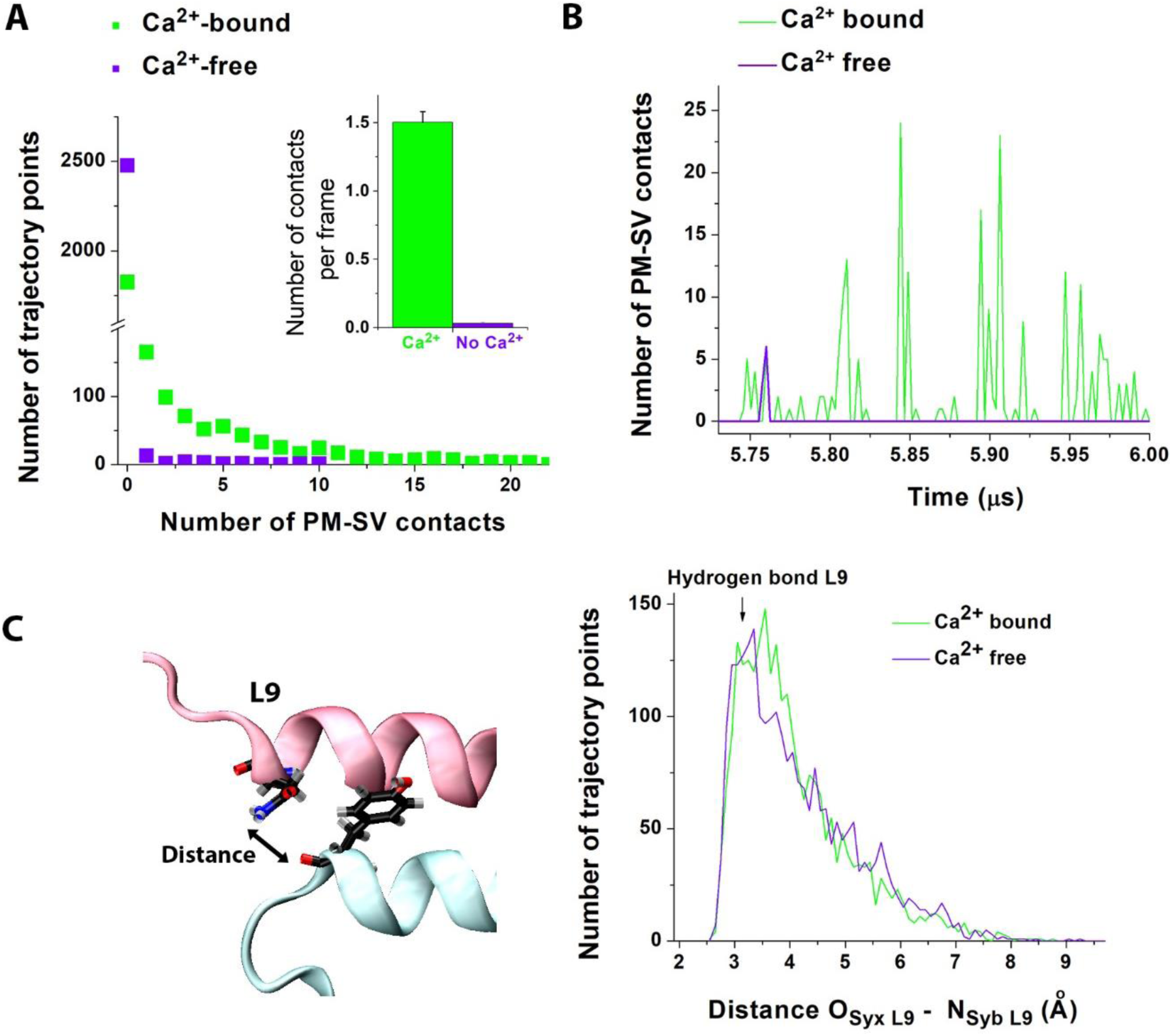
Chelation of Ca^2+^ by Syt1 promotes lipid merging. **A**. The number of contacts between the PM and SV membrane patches over the course of the trajectory is drastically diminished when chelated Ca^2+^ ions are removed from the system. The graph depicts the distribution of the number of VdW contacts observed over the course of the two trajectories. The Ca^2+^ free system mostly shows zero contacts (separated patches), while the system with chelated Ca^2+^ ions shows numerous VdW contacts over hundreds of trajectory points. The inset shows the average number of contacts per frame (p<0.0001). **B**. The number of contacts for Ca^2+^-bound and Ca^2+^-free complex plotted over the final 0.3 μs of the trajectory shows numerous and continuous contacts for the Ca^2+^-bound but not for the Ca^2+^-free system. **C**. SNARE zippering is not altered by the presence of chelated Ca^2+^ in the system. The graph (right) shows the distribution of the distances between the atoms of Syb (nitrogen of N92) and Syx (oxygen of Y257), which form a hydrogen bond when the Layer 9 is fully zippered (shown on the left).

We questioned whether the action of the Ca^2+^C2AB tandem is solely due to its penetration into lipids, or whether the tandem also affects zippering of the Layer 9 of the SNARE complex, either directly, or indirectly via the interactions with lipids. To test this, we quantified zippering of the Layer 9 of the SNARE complex by measuring the distance between its interacting residues (N92 of Syb forming a hydrogen bond with Y257 of Syx, Fig. 11 C). Our results show that zippering of the Layer 9 was not significantly altered by the presence of Ca^2+^ in the complex (Fig. 11 C).

Altogether, these results demonstrate that the force generated by zippering of the SNARE bundle can be sufficient to bring SV and PM in a close proximity but insufficient to overcome the repulsion between the bilayers and to induce the formation of extensive VdW contacts. The additional force needed to merge the two membranes can be generated by the immersion of the Ca^2+^ bound tips of the C2 domains of Syt1 into PM. Syt1 immersed into PM serves as an additional anchor, which also promotes PM curvature, thus bringing PM in a contact with SV. Our results show that this force can trigger the formation of extensive VdW contacts between PM and SV.

## Discussion

We performed the first microsecond-scale all-atom MD simulations of the neuronal Ca^2+^ sensor Syt1 interacting with lipid bilayers and SNARE proteins. The employed time scale of several microseconds allowed us to observe unconstrained conformational transitions, including Syt1 interdomain rotations, the formation of contacts between Syt1 and the SNARE-Cpx bundle, and the immersion of C2 domains of Syt1 into the lipid bilayer.

We started from simulating the interactions of isolated C2 domains with lipid bilayers and subsequently extended our simulations to the C2AB tandem. We next simulated the dynamic interaction of the tandem with the SNARE-Cpx complex and, finally, modeled the prefusion complex comprising the C2AB tandem attached to the SNARE-Cpx bundle positioned between lipid bilayers imitating PM and SV. These simulations produced the model of the prefusion complex, in which Syt1 adopted a conformation with Ca^2+^ bound tips of both domains immersed into PM. The C2B domain was attached to the SNARE bundle, and this enabled the cooperative action of SNARE zippering and Syt1 penetration into PM, which lead to the merging of the PM and SV lipid bilayers *in silico*.

In support of this model, extensive evidence suggested that Ca^2+^ bound tips of both Syt1 domains insert into PM [7, 8, 15], possibly driving the fusion by promoting PM curvature [11, 12, 61]. However, some controversy remained in reconciling these studies with crystallography data [55] supported by computations [48], which suggested that Syt1 adopts a conformation with perpendicularly oriented domains, and therefore the Ca^2+^ bound tips of Syt1 would likely face perpendicular planes. Furthermore, spin labeling studies suggested that membrane-associated Syt1 would adopt a conformation with Ca^2+^ bound tips of its C2 domains facing opposing plains and likely penetrating into opposing bilayers [62]. Our MD simulations resolved this apparent controversy by demonstrating that the interactions of Syt1 with the SNARE complex and with the PM component PIP_2_ collectively promote the conformation of Syt1 in which Ca^2+^ bound tips of both domains can penetrate into PM.

The importance of PIP_2_ in anchoring Syt1 to lipid bilayers has been already established [25, 63-66]. Our MD simulations identified the Syt1 residues interacting with PIP_2_ and determined the 3D configurations for the attachment of each of the C2 modules to lipid bilayers. In particular, the simulations identified three PIP_2_-binding sites for the C2B domain: 1) the polybasic stretch, PB; 2) the basic residues (K366 and K369) of the second Ca^2+^ binding loop, CBL; and 3) the basic residues of the opposite tip (R398 and R399), RR. These residues interchange in forming salt bridges with several PIP_2_ molecules, enabling dynamic and reliable anchoring to a PIP_2_ cluster. In agreement with [63], the simulations showed that such attachment of the C2B domain to a PIP_2_ cluster occurs even in the absence of Ca^2+^. The simulations also showed that upon Ca^2+^ binding the C2B module tilts and its Ca^2+^-bound tip immerses deeper into the bilayer. In this configuration, Ca^2+^ ions form coordination bonds with anionic lipids, but the RR motif is unable to form a salt bridge with PIP_2_ molecules. Thus, both Ca^2+^-free and Ca^2+^-bound forms of the C2B domain anchor to anionic PIP_2_ containing lipids, however the Ca^2+^ -bound form deeper penetrates into the bilayer. These results agree with quantitative thermodynamic studies [64], which observed the Ca^2+^-induced membrane penetration experimentally. Notably, a similar configuration for the attachment of the C2B module to PIP_2_ was demonstrated for the Syt1 analog DOC2B [67], suggesting a general mechanism for the membrane anchoring of C2B domains.

The anchoring mechanism described above can account for the enhanced Ca^2+^ binding observed for the C2B domain of Syt1 in the presence of PIP_2_ [56, 68]. Indeed, the attachment of the CBL motif to PIP_2_ would likely open the Ca^2+^ -binding pocket and allow Ca^2+^ ions to enter the pocket more freely. In addition, the salt bridges formed between PIP_2_ and CBL would neutralize the positive charges of the CBL basic residues and further promote Ca^2+^ entry into the binding pocket. This anchoring mechanisms also agrees with the established role of the PB stretch of the C2B domain [64] and explains the loss of function in Syt1 mutants K366Q [69, 70] and R398Q/R399Q [22].

For the C2A module, our MD simulations revealed only two sites for the interactions with PIP_2_: 1) basic residues of the second Ca^2+^-binding loop (R233 and K236), CBL, and 2) a very short PB stretch containing three only basic residues: K189, K190, and K192. This findings account for the lesser affinity to PIP_2_ observed for the C2A domain versus the C2B domain [71]. The MD simulations also revealed that the Ca^2+^ -binding tip of the C2A domain is rather hydrophobic and forms weak contacts with neutral lipids, unlike the Ca^2+^-binding tip of the C2B domain. The latter result seems to be in an apparent contradiction with liposome binding studies [6], which detected the Ca^2+^-dependent binding of the C2A domain to anionic (POPC:POPS) and PIP_2_-containing liposomes but did not detect a significant attachment of the C2A module to neutral POPC liposomes. However, in agreement with the latter study, our simulations show that the interactions of the C2A domain with the bilayer containing POPS and PIP_2_ are, indeed much stronger than the interactions of the C2A module with POPC. Unfortunately, a direct comparison of MD simulations with the results of *in vitro* experiments is not feasible, since MD approach considers a closed system with all the components positioned in a close proximity to each other. In such a system, even weak affinities between molecules would be detectable, which may be not necessarily the case for *in vitro* assays. It should be noted, however, that the closed system utilized by the MD approach closely imitates the *in vivo* system, whereby Syt1 is attached to SV and positioned in a rather narrow gap between SV and PM. At these conditions, even weak interactions could affect the system dynamics. In particular, even week affinity of the C2A module to neutral lipids would prompt its attachment to SV, as long as PM is not situated in the immediate proximity.

Consistently, the simulations of the C2AB tandem suggested that at a distance from PM the C2A domain would likely have its Ca^2+^ binding tip attached to the SV bilayer, while the C2B domain would likely float in a cytosol. This configuration would be favorable for C2B anchoring to PIP_2_ molecules of PM. Alternatively, the C2B domain could anchor to the SNARE complex.

In agreement with [25], our simulations suggest that the attractions to of the C2B domain to PIP_2_ would be energetically more favorable that the attractions to the SNARE complex. However, besides energetic favorability, other factors should be considered for the molecular system *in vivo*. One important factor is the electrostatic repulsion of SV and PM, which brings up the requirement for molecular anchors. The conformational analysis of Syt1 presented here and in our earlier study [48] suggests that Syt1 could serve only as a relatively short-distance molecular anchor, reaching within a distance of 4-5 nm, since the C2 domains are predominantly coupled and the C2A domain tends to attach to SV. Such a distance range between PM and SV would be reached when the assembly of the SNARE bundle is initiated (Fig. 12 A, 1), and this initial pre-fusion state would be favorable for Syt1 anchoring to t-SNARE.

**Figure 12.**
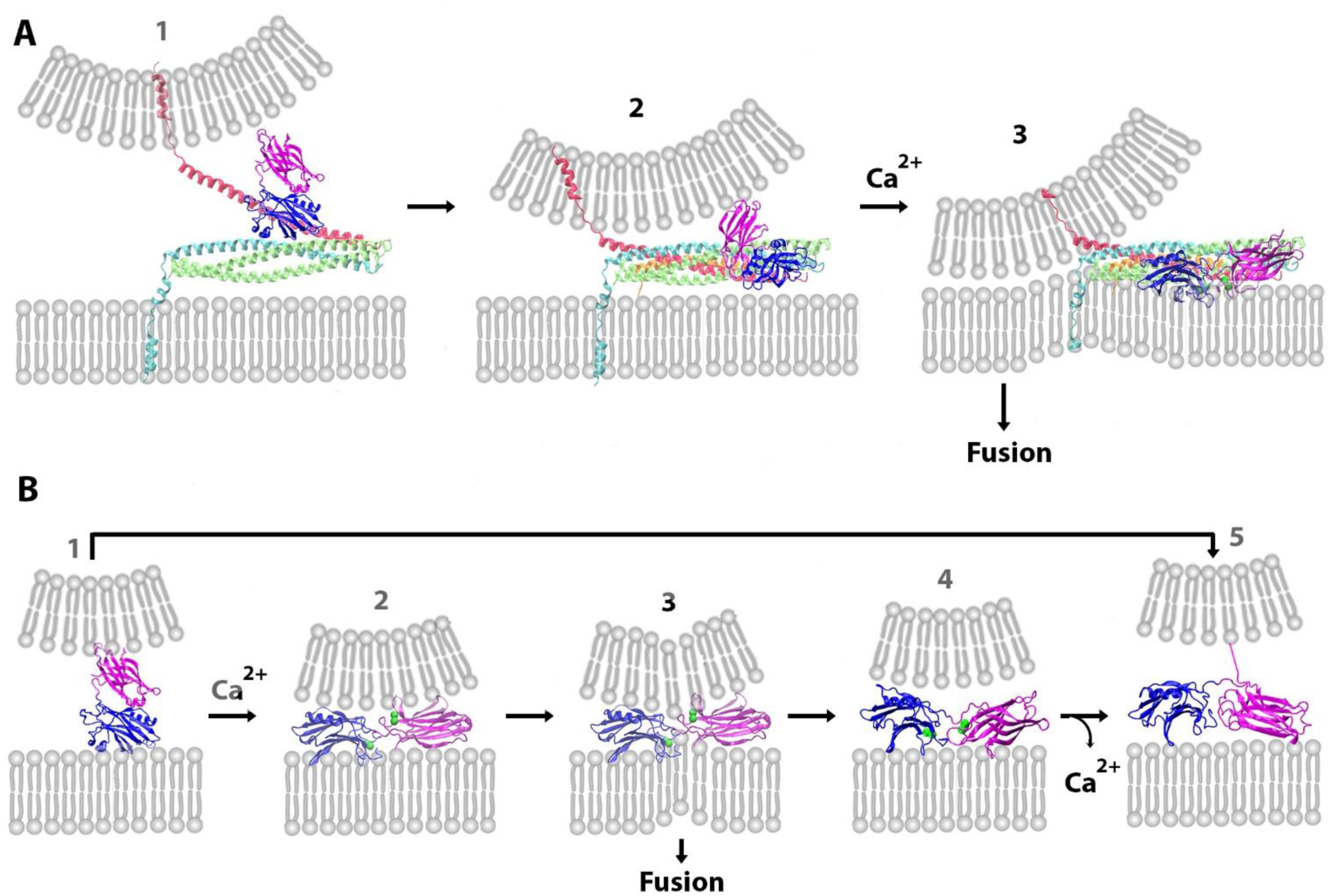
Two hypothetical pathways for the Syt1-mediated fusion. **A**. Synergistic action of the SNARE complex and Syt1. The C2B domain anchors to t-SNARE (1). Subsequently, Syt1 forms an intermediate complex with the SNARE bundle (2). Upon Ca^2+^ binding, both C2 domain immerse into PM, promote PM curvature, and drive fusion synergistically with SNARE zippering (3). **B**. If the SNARE complex is not formed, Syt1 can anchor to PM and drive fusion with a low probability. The C2B domain anchors to PM (1), and upon Ca^2+^ binding the domain tips immerse into opposite bilayers (2), induce lipid budging (3) and can drive lipid merging. However, a conformational transition would likely produce the state with both domain tips penetrating PM (4), which could stabilize (5) and keep SV anchored to PM. The state 1 could also transition to the state 5 directly.

The Syt1-SNARE complex is structurally heterogeneous, and it samples multiple conformational states [57]. Our MD simulations identified three such states with comparable energies for the Syt1-SNARE interactions. The most energetically favorable state (I3, Fig. 8) matched the C2B-SNARE complex identified by the NMR approach [28], and it had the C2B domain anchored to the SNARE complex via its PB stretch. This conformational state is a likely candidate for the initial anchoring of the C2B domain onto the SNARE bundle. However, this conformational state is unlikely to stabilize in the proximity to PM, since PIP_2_ molecules of PM will compete with the SNARE bundle for the attachment to the PB motif of Syt1, and the interactions with PIP_2_ will prevail. Therefore, a subsequent conformational transition would likely occur, leading to a new state of the Syt1-SNARE-Cpx complex. This new conformational state should allow Syt1 to interact with PM and the SNARE bundle simultaneously. Our simulations identified a new conformation of the complex (I2, Fig.8), which appears to be a likely candidate for this intermediate state (Fig. 12 A, 2). The C2B-SNARE interface of this Syt1-SNARE-Cpx complex does not involve the PB motif, thus enabling simultaneous interactions of the C2B domain with the SNARE bundle and PM. Interestingly, this complex has the C2A domain of Syt1 directly interacting with Cpx on the SNARE bundle, thus providing a potential explanation for the role of Cpx in promoting evoked fusion via the interaction with Syt1 [39, 40, 72]. However, structural studies coupled with mutagenesis also propose other pre-fusion states of the Syt1-SNARE-Cpx complex [37], and a sequence of several intermediate states is possible.

Our simulations produced the atomic model of the final pre-fusion state of Ca^2+^Syt1-SNARE-Cpx complex attached to PM (Fig. 12 A.3; Fig. 9). Multiple lines of evidence support this molecular model of the prefusion complex. First, it has an extensive interface between the C2B domain and t-SNARE, which was identified by crystallography [27] and proved stable *in silico* at a microsecond scale at physiological ion concentrations (this study). Second, the mutations disrupting this interface impaired evoked synaptic transmission [37, 47]. Third, the present study showed that the formation of this complex would promote the immersion of both C2 domains of Syt1 into PM, which is thought to be the trigger for evoked Ca^2+^-dependent fusion [8, 11, 15]. Finally, the simulations performed in this study (Fig. 10) showed that this complex would drive lipid merging.

Notably, our simulations elucidated the role of the attachment of the C2B domain of Syt1 to the SNARE bundle. We showed that in the absence of this attachment the C2 domains mutually hinder their immersion into PM, even though they simultaneously anchor to PM. More specifically, the simulations demonstrated that each of the C2 domains in its isolated state would penetrate into PM deeper than within the tandem. This result was surprising, since the bilayer penetration and distortion by one or both C2 domains is thought to drive the synaptic fusion [8, 11, 61]. However, our simulations also demonstrated that the C2B-SNARE attachment uncouples the C2 domains and enables them to immerse into PM deeper. Together with a recent study [32], which demonstrated that Syt1 induces a conformational change in the SNARE complex and promotes its fusion activity, our study supports the cooperative synergistic action of Syt1 and the SNARE complex, suggesting that the Syt1-SNARE interaction mutually promotes Syt1 membrane penetration and SNARE zippering.

Altogether, our simulations favor the scenario for the evoked fusion (Fig. 12 A), in which the C2B domain would anchor onto t-SNARE (Fig. 12 A, 1), and the upon zippering of Syb and attachment of Cpx would transition through intermediate state(s), likely involving the interaction between Syt1 and Cpx (Fig. 12 A, 2). Upon Ca^2+^ binding, the complex would transition to its final prefusion state (Fig. 12 A,3), and the immersion of the C2 domains into PM would drive membrane curvature followed by fusion.

Syt1 could also anchor to PM independently of the SNARE complex (Fig. 12 B). Being a transmembrane protein attached to SV and having a high affinity to the PM component PIP_2_, Syt1 would serve as an efficient anchor for docking SVs to PM [24, 25]. In addition, our simulations identified a potentially fusogenic state of Ca^2+^Syt1, which has the Ca^2+^ bound tip of the C2A domain attached to the SV bilayer, and the Ca^2+^ bound tip of the C2B domain immersed into the PM bilayer (Fig. 12 B, 3; Fig. 7). This finding agrees with a spin labeling study [10], which suggested that Syt1 could bridge membranes by penetrating the opposing bilayers with the Ca^2+^-bound tips of its C2A and C2B domains. Our simulations showed that such state of the C2AB tandem would promote lipid bulging, and this could potentially lead to fusion. However, the simulations also showed that unless the lipid merging occurs rapidly, such state of the C2AB tandem would not stabilize, since Syt1 would likely undergo a conformational transition so that the Ca^2+^ bound tip of the C2A domain would attach to PM (Fig. 12 B, 4). Therefore, this pathway could serve to promote the SV-PM docking and possibly mediate low-probability fusion, such as asynchronous or spontaneous transmitter release [73]. This model (Fig. 12) agrees with the finding that different conformational states of Syt1 control different components of synaptic fusion [15].

## Methods

### System setup

The POPC lipid bilayers were generated using VMD. The initial structure for the POPC:POPS:PIP_2_ bilayer (75:20:5) [74] was kindly provided by Dr. J. Wereszczynski (Illinois Institute of Technology). In all the systems, the lipid bilayers were positioned in the XY plain.

## Acknowledgements

This study was supported by the NIMH grant R01 MH099557. MD simulations were performed at the Anton supercomputer (D.E. Show Research and Pittsburg Supercomputer Center) and at XSEDE resources (Stampede supercomputer at TACC). We thank Dr. J. Wereszczynski for kindly sharing the model of POPC:POPS:PIP_2_ membrane.

