## Supplemental Table 1 and supplemetal figures 1-12 for "SNARE Complex Alters the Interactions of the Ca^2+^ sensor Synaptotagmin 1 with Lipid Bilayers"

### Supporting Material

***Table S1*. The summary of the performed MD simulations.**

| System | Size (atoms), including water molecules and ions | Trajectory length (μs) | Figures showing the results |
|---|---|---|---|
| C2B – POPC | 50362 | 3.2 | 1, S1 |
| $Ca^{2+}$C2B – POPC | 49118 | 4.0 | 1, S1 |
| C2B – POPC:POPS:$PIP_2$ | 49820 | 3.9 | 1, S2 |
| $Ca^{2+}$C2B - POPC:POPS:$PIP_2$ | 49821 | 4.4 | 1, S2 |
| $Ca^{2+}$ C2B - SNARE-Cpx – POPC:POPS:$PIP_2$ | 235563 | 3.6 | 2, S3 |
| C2A – POPC | 50037 | 3.8 | 3, S4 |
| $Ca^{2+}$C2A – POPC | 50040 | 3.7 | 3, S4 |
| C2A – (POPC:POPS:$PIP_2$ ) | 49821 | 3.9 | 3, S5 |
| $Ca^{2+}$C2A – (POPC-POPS-$PIP_2$) | 49824 | 3.9 | 3, S5 |
| C2AB – POPC | 108340 | 4.7 | 4, S6 |
| $Ca^{2+}$C2AB – POPC | 108328 | 4.9 | 4, S6 |
| C2AB – POPC:POPS:$PIP_2$ | 107011 | 5.1 | 5, S7 |
| $Ca^{2+}$C2AB - POPC:POPS:$PIP_2$ | 106807 | 5.0 | 6, S7 |
| $Ca^{2+}$C2AB – POPC:POPS-$PIP_2$/POPC | 60505 | 6.6 | 7 |
| $Ca^{2+}$C2AB – SNARE - Cpx<br>Starting from X-ray (Zhou et al., 2015) | 231560 | 9.0 | S8 |
| I1: C2AB – SNARE - Cpx | 277270 | 8.8 | 8, S9, S10 |
| I2: C2AB – SNARE - Cpx | 224739 | 7.0 | 8, S9, S10 |

| | | | |
|---|---|---|---|
| I3: C2AB – SNARE - Cpx | 278775 | 6.1 | 8, S9, S10 |
| I4: C2AB – SNARE - Cpx | 246451 | 2.7 | S9, S10 |
| $Ca^{2+}$C2AB – SNARE - Cpx - POPC:POPS-$PIP_2$ | 175447 | 8.4 | 9 |
| POPC - $Ca^{2+}$C2AB-SNARE-Cpx - POPC:POPS:$PIP_2$ | 361130 | 6.0 | 10, S11 |
| POPC – C2AB–SNARE-Cpx - POPC:POPS-$PIP_2$ | 347329 | 6.0 | 10, S12 |

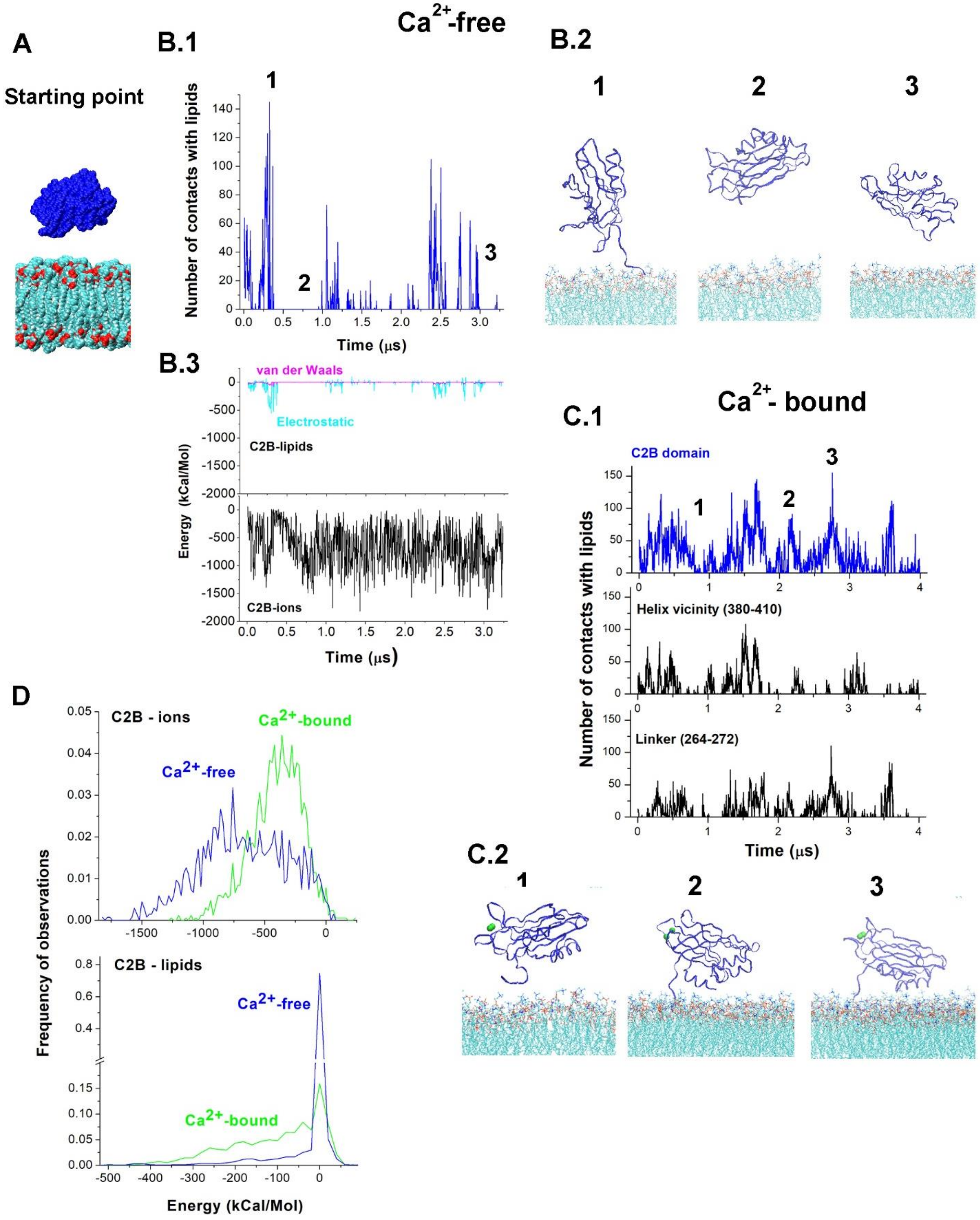


***Figure S1.*** Interactions of the C2B domain with the neutral lipid bilayer. **A.** The starting configuration: a VdW representation of the C2B domain positioned over the POPC homogeneous lipid bilayer. **B.** The MD trajectory shows that the $Ca^{2+}$ -free form of the C2B domain is largely detached from the bilayer and has a preference for water/ion environment. **B.1.** The number of VdW contacts over the course of the MD trajectory. The time-points 1,2, and 3 correspond to the states shown in B.2. **B.2.** Representative snapshots from the MD trajectory. **B.3.** The energy components of the protein-lipid interactions versus protein-ion interactions show that protein-ion interactions would prevail. **C.** The MD trajectory of the $Ca^{2+}$C2B module shows numerous contacts between the protein and the bilayer. **C.1.** $Ca^{2+}$C2B-lipid **c**ontacts over the course of the MD trajectory (blue) largely reflect the interactions of the α-helix and the adjacent β-sheets (residues 380-410) or the interdomain linker (residues 264-272) with the bilayer. The time-points 1,2, and 3 correspond to the states shown in C.2. **C.2.** Representative snapshots from the MD trajectory of the $Ca^{2+}$C2B module. Note the proximity of the α-helix to the bilayer and $Ca^{2+}$-binding loops facing away from the bilayer. Green spheres – $Ca^{2+}$. **D**. The distributions of the energies of protein-lipid and protein-ion interactions for the $Ca^{2+}$-free and $Ca^{2+}$ bound forms of the C2B domain along respective trajectories. Note that the energies of the protein-ion interactions are shifted towards more negative values for the $Ca^{2+}$-free form ($p<0.001$). In contrast, the energies of the protein-lipid interactions are shifted towards more negative values for $Ca^{2+}$C2B ($p<0.001$).

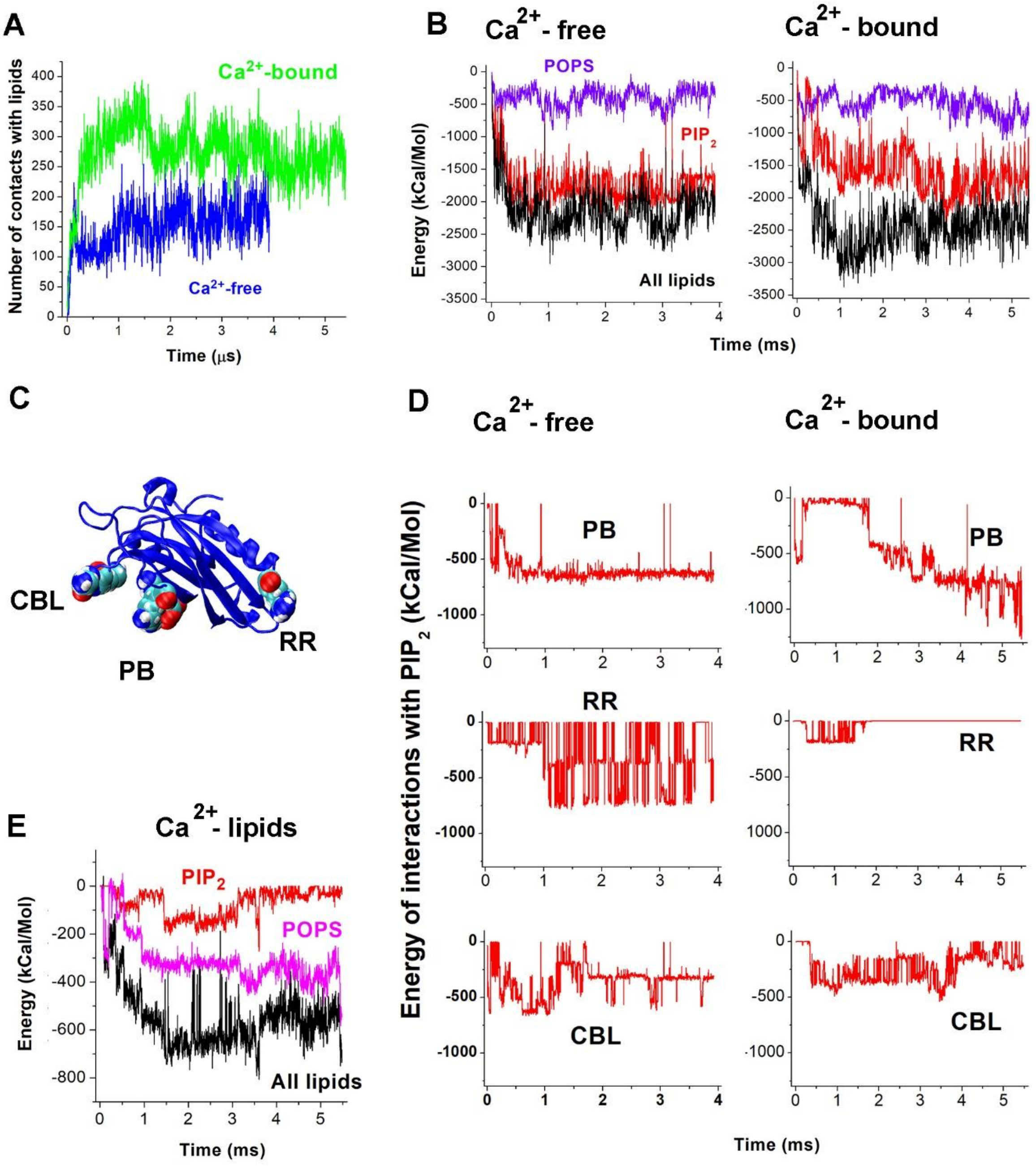


***Figure S2.*** The interactions of the C2B domain with the POPC:POPS:$PIP_2$ lipid bilayer. **A.** The number of VdW contacts between the $Ca^{2+}$-free (blue) or the $Ca^{2+}$-bound (green) C2B module and the lipid bilayer over the course of the MD trajectory. Note an increase in the number of contacts during the first microsecond followed by a plateau. The plateau level is higher for the $Ca^{2+}$-bound C2B form, showing more extensive contacts with the bilayer. **B.** The energy of the C2B-lipid (black lines) interactions is largely defined by the interactions between the C2B module and $PIP_2$ (red lines), although the interactions with POPS (purple) provide a modest contribution to the overall enthalpy, especially for the $Ca^{2+}$ bound form (right). **C.** Three structural elements of the C2B domain defining its interactions with $PIP_2$: polybasic stretch K321-K327 (PB); basic residues of the 2nd $Ca^{2+}$-binding loop, K366 and K369 (CBL); and two arginines at the opposite tip, R398-399 (RR). **D.** Energy of the interactions of the three motifs of the C2B domain (PB, RR, and CBL) with $PIP_2$. Note that all the three elements dynamically interact with $PIP_2$ for the $Ca^{2+}$-free C2B module (left panels). In contrast, for the $Ca^{2+}$ bound C2B module the energy of RR-PIP2 interactions becomes zero after the initial 2 μs of the simulations, showing the lack of binding (right middle panel). **E.** The contribution of $Ca^{2+}$ ions into the interactions of $Ca^{2+}$C2B with lipids. The energy of $Ca^{2+}$-lipid interactions is largely defined by the electrostatic attraction of $Ca^{2+}$ ions to POPS (magenta).

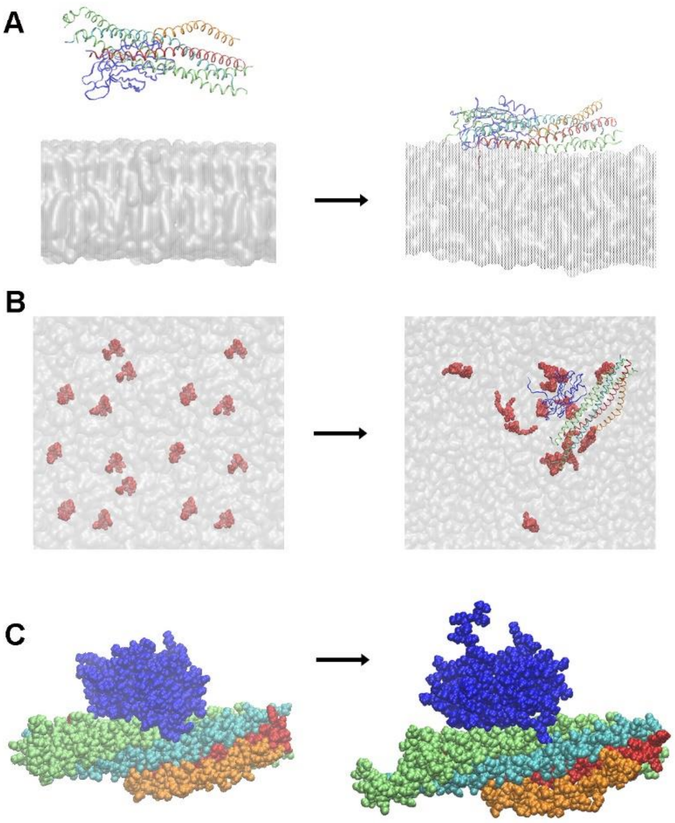


***Figure S3.*** Changes in the $Ca^{2+}$C2B-SNARE-Cpx complex on the POPC:POPS:$PIP_2$ bilayer over the course of the trajectory. **A.** The initial configuration (left) has the complex detached from the membrane, while over the course of the simulation the complex attached tightly to the bilayer (right). **B.** Rearrangement of $PIP_2$ molecules. The initial membrane patch (left) had evenly distributed $PIP_2$ (red, VdW representation). Over the course of the MD run, the $PIP_2$ molecules aggregated near the protein complex (right). **C.** The overall configuration of the protein complex did not substantially change over the course of the simulation. VdW representations of the initial (left) and final (right) show similar structures. Red: Syb, cyan: Syx, lime: SNAP25, orange: Cpx, blue: C2B. Note a tight interaction between C2B and SNAP25.

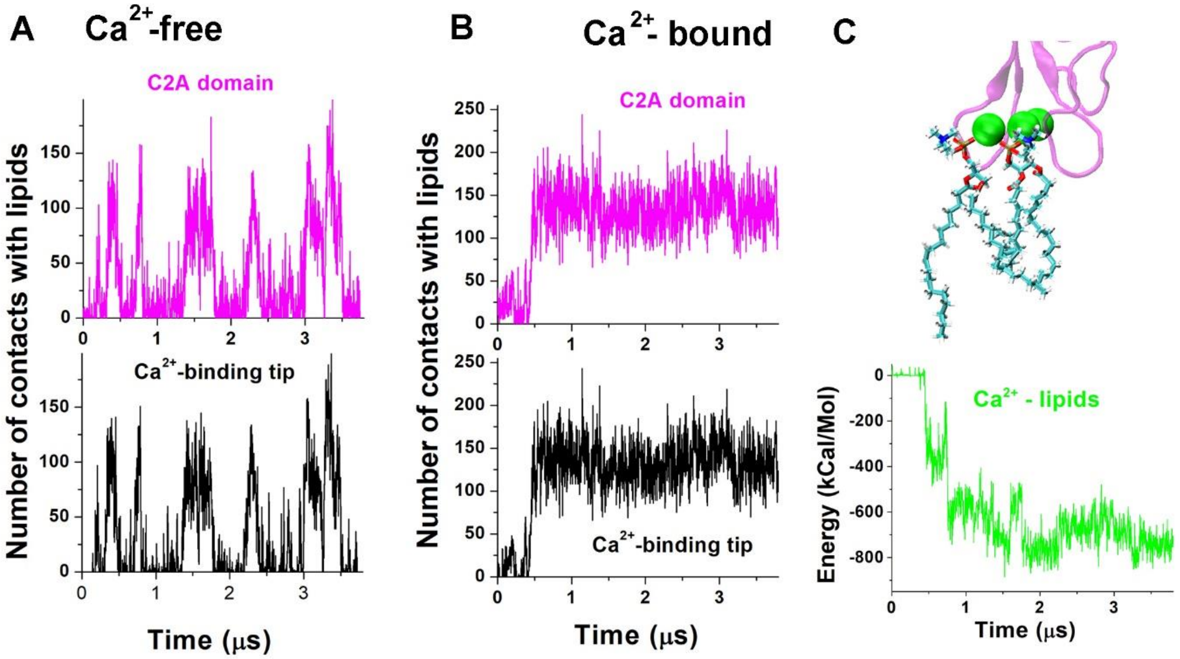


***Figure S4.*** Interactions of the C2A domain with neutral lipids (POPC). **A, B.** C2A-lipid **c**ontacts along the MD trajectory (magenta) closely matches the contacts between the three loops of its $Ca^{2+}$-binding tip and the bilayer. **C.** $Ca^{2+}$ ions form coordination bonds with oxygen atoms of phosphate groups of POPC molecules (top), while the energy of $Ca^{2+}$ -POPS interactions decreases over the course of the trajectory and reaches a plateau after 2 μs (bottom).

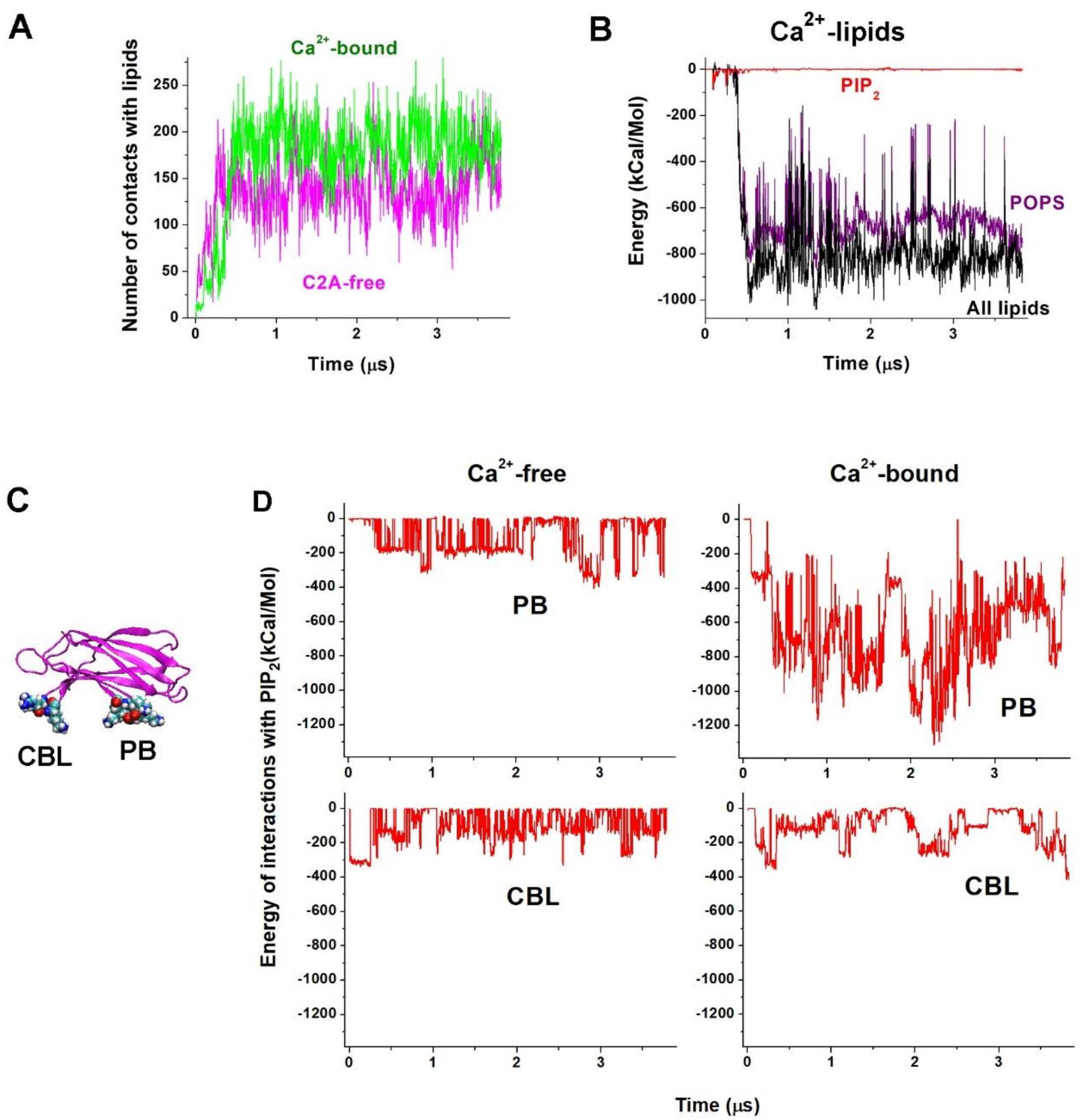


***Figure S5.*** Interactions of the C2A domain with the lipid bilayer POPC:POPS:$PIP_2$.**A.** The protein rapidly attached to bilayer, and the attachment (the number of VdW contacts) is more extensive for the $Ca^{2+}$-bound form of the C2A module. **B.** $Ca^{2+}$ ions contribute to the interactions with the bilayer predominantly via their interactions with the anionic lipids POPS. **C.** Two anchors of the C2A module forming salt bridges with $PIP_2$ : PB (K189-193) and CBL (R233 and K236). **D.** The energy of the interactions between $PIP_2$ and the anchors PB and CBL along the respective trajectories.

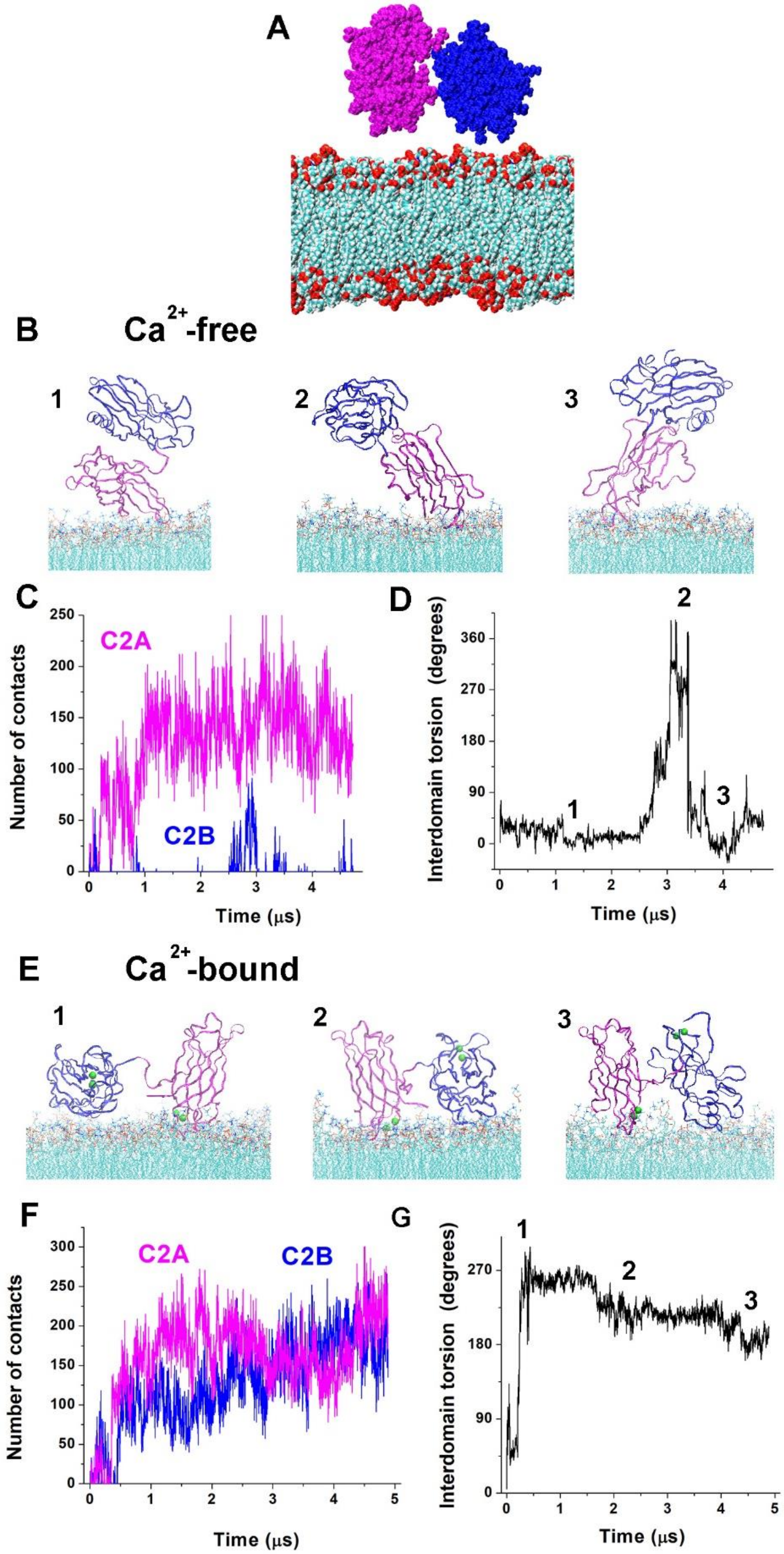


***Figure S6.*** Trajectories for the C2AB tandem interacting with the neutral lipid bilayer (POPC). **A.** The initial state: C2 domains are not tightly coupled and detached from the bilayer. **B.** The $Ca^{2+}$-free C2AB tandem transitions trough multiple conformational states, all of which have the C2B domain detached from the bilayer and the C2A domain attached to the bilayer via its $Ca^{2+}$-binding tip. **C.** The number of C2A-POPC contacts increases along the MD trajectory and reaches a plateau, while the C2B-POPC contacts are infrequently. **D.** Changes in the interdomain torsion angle (residues G305-E350-N154-M173 (Bykhovskaia, 2015)) along the trajectory showing the rotation of C2 domains. Time points 1,2,3 correspond to the conformational states in the panel B. **E.** Conformational states of $Ca^{2+}$C2AB: both domains are attached to the bilayer. **F.** The number of contacts increases along the MD trajectory for both domains. **G.** Changes in the interdomain torsion angle of $Ca^{2+}$CAB showing the domain rotations, with the time points 1,2,3 corresponding to the conformational states in the panel E.

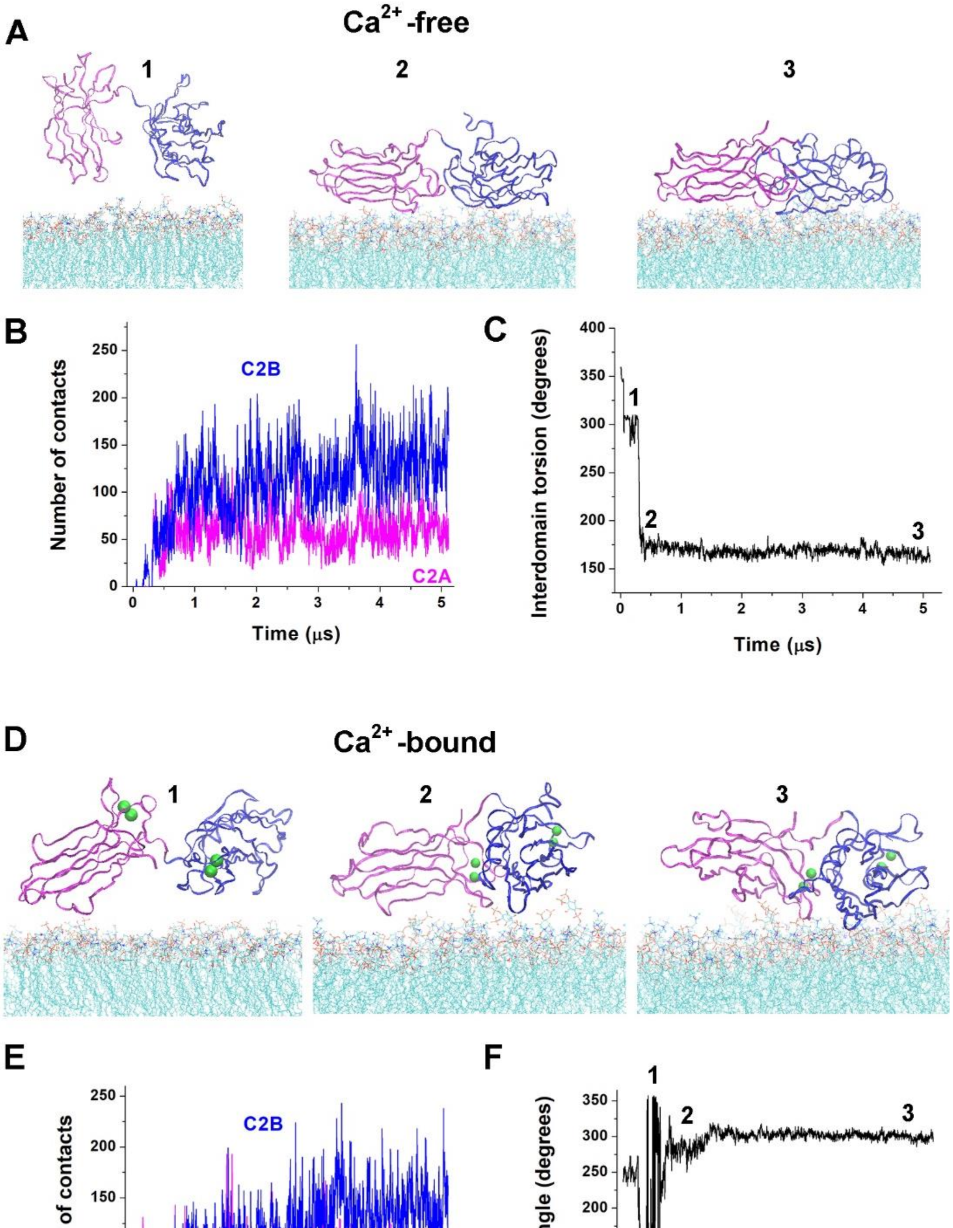


***Figure S7.*** The trajectory of the C2AB tandem interacting with POPC:POPS:$PIP_2$. **A.** Time-points along the trajectory of the $Ca^{2+}$-free C2AB tandem show the protein attachment to the bialyer. **B.** VdW contacts between each of the C2 domains and the bilayer increase along the MD trajectory and reach a plateau. **C.** The interdomain torsion angle shows the rotation of C2 domains within the initial 500 ns. Time points 1,2,3 correspond to the conformational states in the panel A. **D.** Conformational states of $Ca^{2+}$C2AB along the trajectory. **E.** Contacts with the bilayer increase along the MD trajectory for eachdomains and reach a plateu. **F.** The interdomain torsion angle of $Ca^{2+}$CAB shows multiple domain rotations within the initial 1 μs of the trajectory. The time points 1,2,3 correspond to the conformational states in the panel D.

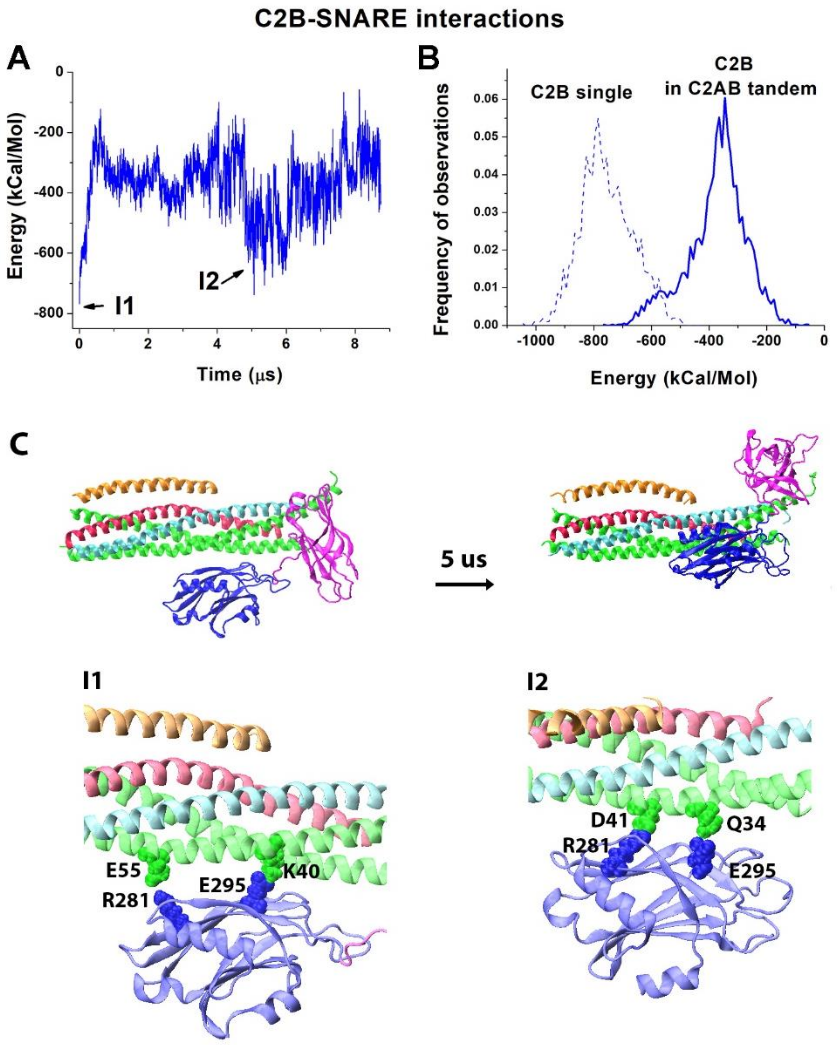


***Figure S8.*** The initial trajectory of the CAB-SNARE-Cpx complex suggests additional conformational state(s). **A.** The energy of the C2B-SNARE interactions rapidly increases from the initial state (I1), plateus, and the decends into a local minimum (I2). The energy of the C2B-SNARE interactions is signficantly ($p<0.001$) lower for the isolated C2B domain compared to the C2B domain within the C2AB tandem, suggesting that the C2A domain destabilizes the complex in this conformation. **C.** The conformtaional transition of the complex from I1 to I2 C2B-SNARE interface. The initial I1 interface is characterized by two salt bridges between R281 and E295 of the C2B domain (blue) and E55 and K40 of SNAP25 (green). In the I2 complex, the C2B domain is shifted along the SNARE bundle towards its membrane-distal end by approximately 3 layers. In the I2 complex, the same residues of the C2B domain (R281 and E295) form salt bridges with D41 and Q34 of SNAP25.  Red: Syb; cyan: Syx; green: SNAP25; orange: Cpx.

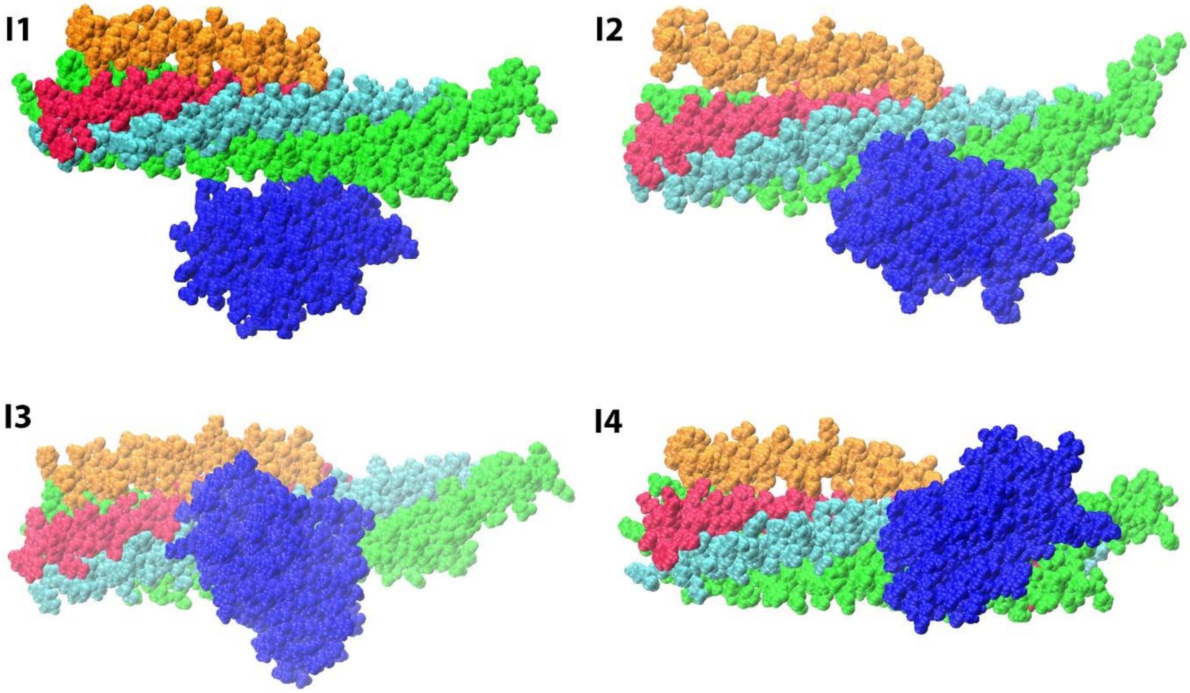


***Figure S9***. Four initial configurations for the attachment of the C2B domain to the SNARE-Cpx bundle. The protein complexes are shown in the VdW representation. Red: Syb; cyan: Syx; green: SNAP25; orange: Cpx; blue: C2B domain of Syt1.

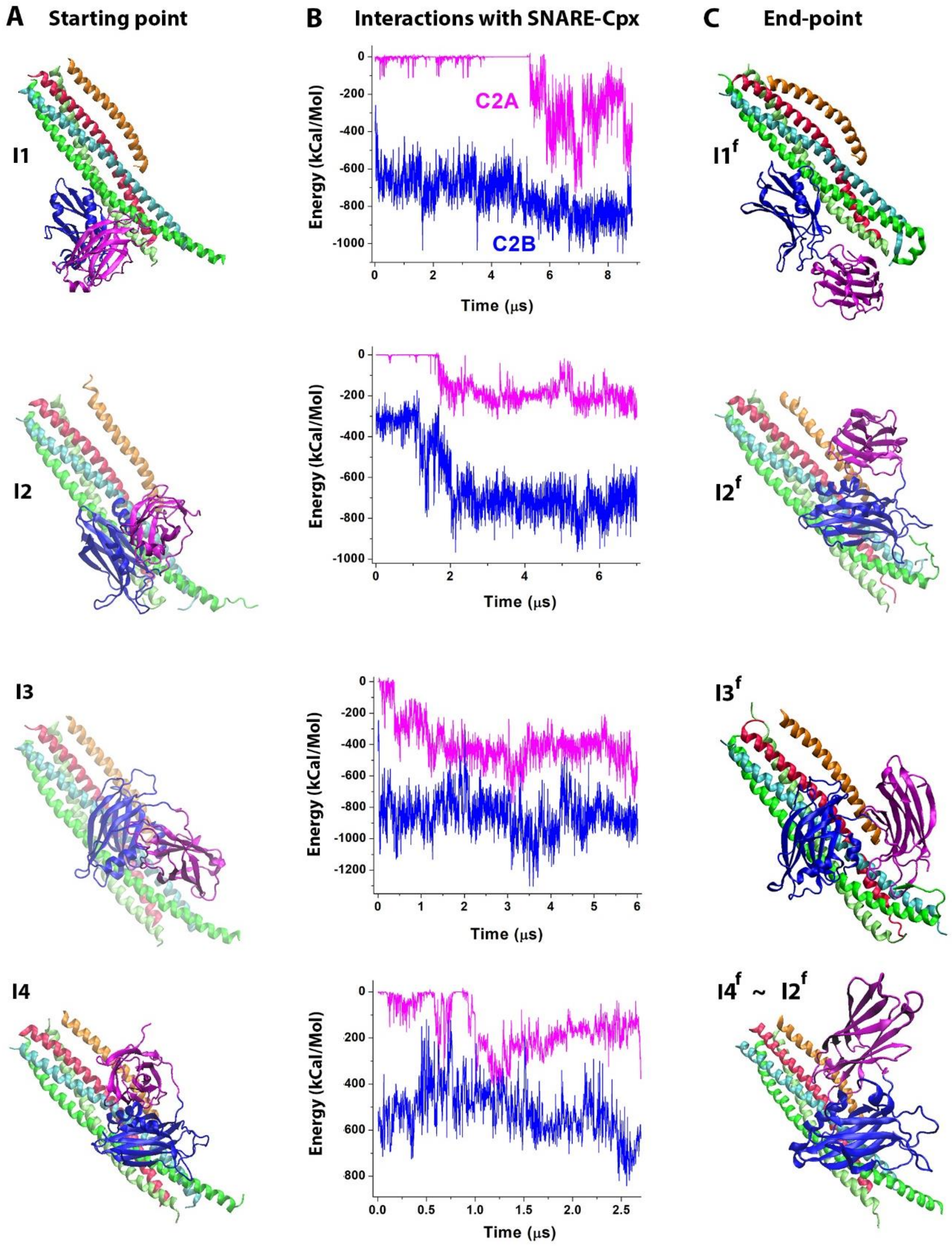


***Figure S10.*** The MD trajectories of the C2AB-SNARE-Cpx complex. **A.** Four initial approximations: I1 – I4. **B.** The energy profiles for the interactions of the C2 domains with the SNARE-Cpx bundle. **C.** The endpoints of the four trajectories. Note that the end-point $I4^f$ has topological similariteies with the end-point $I2^f$, including the orientation of the C2B domain on the SNARE bundle and the attachment of the $Ca^{2+}$-binding tip of the C2A domain to Cpx.

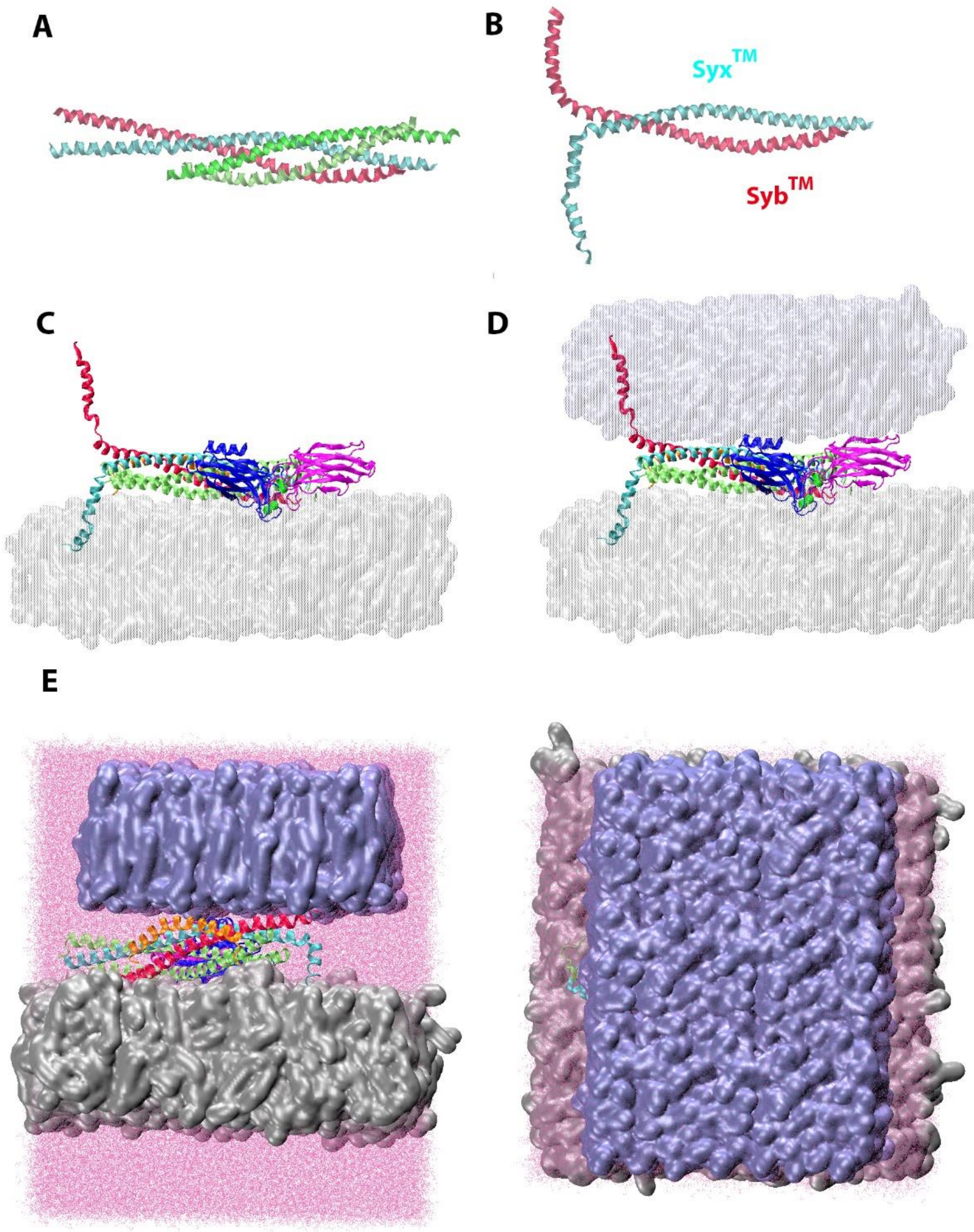


***Figure S11.*** Building the model of the pre-fusion complex. **A** The structure of the SNARE complex with transmembrane domains, 3IPD, (Stein, Weber, Wahl, & Jahn, 2009). Red: Syb$^{TM}$, cyan: Syx$^{TM}$, green: SNAP25. **B.** The Syx$^{TM}$-Syb$^{TM}$ complex, constructed from the 3IPD SNARE complex by removing SNAP25 and bending the transmembrane domains. **C.** The Syx and Syb proteins in the $Ca^{2+}$I1-PM complex were substituted by Syx$^{TM}$ and Syb$^{TM}$, respectively. **D.** The POPC membrane patch was added on the top of the protein complex. **E.** Two perpendicular views of the periodic cell, showing the sizes of the membane pathcs and water molecules. The POPC:POPS:$PIP_2$ patch – silver, surface representation; the POPC patch – ice blue, surface representation; water: pink, dot reprresentation.

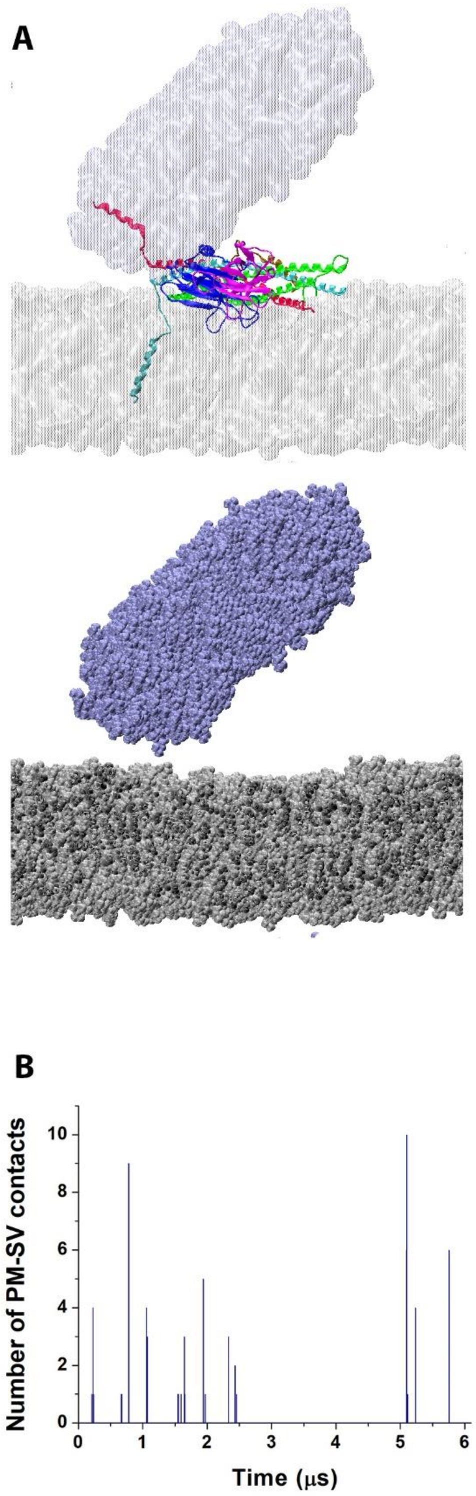


***Figure S12***. In the absence of $Ca^{2+}$, the C2AB-SNARE-complex drives the two membrane patches together, but not to the point where extensive VdW contacts are formed. **A.** The end-point of the trajectory. Two representations show the C2AB-SNARE-Cpx complex between the bilayers, which approached each other but did not make VdW contacts. The top view shows the protein complex between the bilayers. The bottom view shows the bilayers in VdW representation, with the protein complex being removed for claritty. **B.** The number of VdW contact between the bilayers along the trajectory. Note that the contacts are only formed occationally, and they are not numerous.
